# Genome-wide investigation of Cytochrome P450 superfamily of *Aquilaria agallocha*: association with terpenoids and phenylpropanoids biosynthesis

**DOI:** 10.1101/2022.09.26.509443

**Authors:** Ankur Das, Khaleda Begum, Suraiya Akhtar, Raja Ahmed, Phatik Tamuli, Ram Kulkarni, Sofia Banu

## Abstract

P450 superfamily (CYPs) has been known as contributors to the metabolites’ diversity and their promiscuous nature has led to the flexibility in substrate specificity and functional diversity. Current study was designed to investigate CYPs in the genome of an agarwood producing plant species named Aquilaria agallocha. Agarwood, the resinous fragrant wood with numerous phytochemicals, produced when an Aquilaria plant respond to wound and microbial infection. These chemicals are of great interest to industries ascribing it a high economic value. However, the pathways for the biosynthesis of these metabolites have not been studied in context of Aquilaria CYPs. We identified 136 A. agallocha CYP proteins from the genome, characterized and classified them into 8 clans and 38 families. Functional analysis unveiled their participation in terpenoids, phenolics, flavonoids and other valuable metabolites biosynthesis. Conserved motifs were detected and evolutionary analysis revealed duplicated and orthologous pairs. Potential members for the biosynthesis of sesquiterpenoids and phenylpropanoids reported in Aquilaria and agarwood were elucidated and validated through expression profiles in stress induced callus tissues and infected Aquilaria tress. This study provides a strong foundation for biochemical characterization of Aquilaria CYPs which will aid heterologous production of valuable phytochemicals and untangle molecular mechanism of agarwood formation.

## 1. Introduction

Cytochrome P450s (CYPs) belong to the largest and diverse protein family ubiquitous to all living beings (Nelson and Werck-Reichhart 2011). Owing to its versatility of substrate catalysis and involvement in diverse metabolic pathways, the superfamily has remained an attraction to many researchers, and with due course, increasing evidences have confirmed their relevance in all forms of life (Guengerich et al. 2021, Liu et al. 2020, Peng et al. 2017). CYPs catalyse a wide-range of chemical reactions including hydroxylation, oxidation, epoxidation, reduction, decarboxylation, isomerization, C-C cleavage, ring extension including stereospecific catalysis and so on (Mizutani et al. 2010). In angiosperms, they participate in metabolism of xenobiotics (pesticides and pollutants) into nontoxic (Werck-Reichhart and Feyereisen 2000), and secondary metabolism such as biosynthesis of terpenes, phytohormones, sterols, and lignin (Xu et al. 2015). CYPs utilize heme-thiolate for molecular oxygen reduction and their conserved residues, viz., heme-binding motif (FXXGXRXCXG), K-helix (EXXR), I-helix (AGXD/ET), and PXRX motif for the catalysis (Graham-Lotence et al. 1995, Denisov et al. 2005).

Nearly 200,000 CYPs from various plant species have been reported till date (Nelson 2018) and are periodically updated and maintained in the Cytochrome P450 database (Nelson et al. 2009). Genome level identification of CYPS have been carried out in numerous plants including *Vitis vinifera* (Jiu et al. 2020); *Oryza sativa* (Wei at al. 2018); *Ricinus communis* (Kumar et al. 2014); *Morus notabilis* (Ma et al. 2014); *Nicotiana tabacum* (Xie et al. 2013); *Nelumbo nucifera* (Nelson 2013); *Linum usitatissimum* (Babu et al. 2013); *Glycine max* (Guttikonda et al. 2010); *Populus trichocarpa* (Nelson et al. 2008), *Medicago truncatula* (Li et al. 2007), and *Arabidopsis thaliana* (Xu et al. 2001). Based on phylogenetic position and sequence similarity, the CYPs are divided into 10 clans i.e., single-family clans (51, 74, 97, 710, 711, and 727) and multiple-family clans (71, 72, 85, and 86) (Nelson et al. 1996). In plants, members of CYP71 clan have been categorized into A-type, and rest into non-A type (Paquette et al. 2000, Nelson 1999). In spite of quite a number of sequence identified, very less members has been biochemically characterized (Wang et al. 2021). The members of CYP71 clan are involved in terpenoids, amino acid derivatives, hormone precursors, and alkaloids metabolism (Nelson and Werck-Reichhart 2011). The families of CYP85 clan in triterpenoid biosynthesis (Misra et al. 2017, Moses et al. 2015), CYP72 clan in phytohormones, fatty acids, cytokinins, and terpenoids biosynthesis (Nelson and Werck-Reichhart 2011), and CYP86 clan in fatty acid metabolism (Pinot and Beisson 2011). The CYP73, CYP84, and CYP98 clan members are involved in flavonoids, suberin, lignin, coumarins, and polyphenol biosynthesis (Mizutani and Ohta. 2010), whereas, CYP710 and CYP51 clan members participate in biosynthesis of sterol (Lepesheva and Waterman 2007, Morikawa et al. 2009). Similarly, the CYP711 and CYP97 members involved in carotenoid (Booker et al. 2005, Kim et al. 2010) and CYP74 in fatty acid metabolism (Hughes et al. 2009). CYPs also play role during biotic and/or abiotic stresses such as temperature (Qin et al. 2008), drought (Tamiru et al. 2015), heavy metal (Rai et al. 2015), salt (Wang et al. 2017), insects and pest infection (Schuler et al. 1996, Li et al. 2010). Furthermore, biotic agents such as insects, microbes, and weeds are also responsible for inducing expression of *CYPs* associated with defence compounds and secondary metabolites biosynthesis (Wasternack et al. 2016, Butrón et al. 2010, Bari et al. 2009).

*Aquilaria* spp. are known for their association with a rare phenomenon called agarwood. As a result of wound and microbial infection, *Aquilaria* tress start to deposit fragrant metabolites which eventually turns the softwood into dark resinous agarwood (Mohamed et al. 2014, Subasinghe and Hettiarachchi. 2013). It has been in Chinese medicine for long as flatulence and pain reliever, stomachic and tranquilizer (Liu et al. 2017), and fascinated various pharmaceutical and perfumery industries (Liu et al. 2013). With a global market up to US$ 8 billion, agarwood has become a major economic attraction (Tan et al. 2019). The chemical constituents that impart agarwood and *Aquilaria* plants its value are mostly terpenoids, flavonoids, phenolics and phenylethyl chromones (Wang et al. 2018). Among these, sesquiterpenoids, phenyl chromones and their derivatives have been most commonly isolated and studied (Gao et al. 2019, Ahmaed et al. 2017). Metabolites profiling of different grades of agarwood has shown a direct correlation of its quality with the chemical constituents (Islam et al. 2022, Liu et al. 2017), which are hydroxylated and oxygenated products viz. sesquiterpenoids, 2-(2-phenylethyl) chromones (PECs) and few conjugates (Li et al. 2021). It is believed that the unique fragrance that agarwood exhibits is because of these metabolites (Tan et al. 2019). Detection of such unique chemical profile indicates CYPs action of hydroxylation and oxidation of sesquiterpenes (Chappell et al. 2010) and precursors of phenylethyl chromones (Wang et al. 2022) during agarwood formation in *Aquilaria* plant species.

Considering the evidences of CYPs in other species and no documentation of *Aquilaria* CYP450s till date, we assume that CYPs in *Aquilaria*, may also be involved in the biosynthesis of the natural products in agarwood. We intended to identify and functionally classify CYPs in *Aquilaria agallocha*, which would provide opportunities to explore their functional biology in *Aquilaria*, and give insight into molecular mechanism for biosynthesis of agarwood chemicals and other economically important metabolites.

## 2. Methods and materials

### 2.1. Plant materials, callus establishment and MeJ treatment

Wood tissue of Agarwood producing (infected) and healthy (non-infected) *Aquilaria* tress were obtained from Hoollongapar Gibbon Sanctuary, Jorhat, India as per methodology described by Islam et al. 2020. Samples were transported in liquid nitrogen and stored at −80°C. For tissue culture, leaves were disinfected using sodium hypochlorite solution and cleansed with sterile water. Subsequently, cut into small pieces and aseptically inoculated in Murashige and Skoog (MS) medium (pH 6.8) with 3% (w/v) sucrose enriched with 2, 4-Dichlorophenoxyacetic acid (2, 4-D) at 6 mg/L, and kinetin at 2 mg/L, and incubated at 25 °C in dark (Okudera et al. 2009). Well grown callus was used for cell suspension culture, and after 5 days of incubation at 130 rpm, methyl jasmonate (MeJ) dissolved in DMSO was added at concentration of 0.1 mM (Kumeta and Ito 2010). After 48 hours, the cells were harvested through filtration and immediately frozen in liquid nitrogen and stored at −80°C.

### 2.2. Identification of *CP450s* in the *Aquilaria* genome

The protein sequences were obtained from our previous annotation project (Das et al. 2021). P450 domain (PF00067) of 149 plant species was downloaded to generate HMM profile utilizing HMMER 3.0 (Potter et al. 2018). To identify *A. agallocha* CYPs (*AaCYPs*), the sequences were subjected to hmmsearch against the P450 profile. Thereafter, the short and redundant sequences were omitted, and P450 domains were confirmed with ScanProsite server (Sigrist et al. 2002) and Pfam database. The candidate CYPs were then blast searched to Uniprot database to reconfirm their identity and find closest match. Number of transmembrane regions were predicted with TMHMM (Möller et al. 2001), while the molecular weights, localization and theoretical PI were calculated with ExPASy server.

### 2.3. Phylogenetic relation, gene structures and conserved motifs

For phylogenetic analysis, reviewed CYPs of *Arabidopsis thaliana* from UniProt database were obtained followed by sequence alignment and a neighbor joining tree construction with MEGA 7.0 (Kumar et al. 2018). Gene structure display server 2.0 (Hu et al. 2015) was employed to map the intron and exon boundaries from genomic DNA and cDNA sequences. The cis-regulatory elements (CREs) in the upstream regions of *AaCYPs* were identified from plant CARE database (Lescot et al. 2002). Conserved motifs were determined with MEME Suite as per parameters mentioned by Liu et al. 2018.

### 2.4. Synteny and duplication events

The genomic and protein sequences of *Vitis vinifera, Arabidopsis thaliana, Solanum lycopersicum*, and *Glycine max* were obtained from Phytozome v13, and syntenic relations were analysed with MCScanX (Wang et al. 2012) and presented with TBtools (Chen et al. 2020). Duplicated *AaCYPs* genes were identified through blastp with cut-off identity >75% and coverage >75% of query length. Two *AaCYPs* if located >100 kb distance were considered segmental duplication, while genes located <100 kb distance within same scaffold were considered tandem duplication (Islam et al. 2019; Lopez-Ortiz et al. 2019). Synonymous (Ks) as well as non-synonymous (Ka) rate of substitution was evaluated with KaKs_calculator (Wang et al. 2010) and diversion time as per Moniz de Sa and Drouin 1996.

### 2.5. Mapping, expression and pathway enrichment analysis

AaCYPs were mapped with Kyoto Encyclopedia of Genes and Genomes (KEGG) and Gene ontology (GO) database for functional prediction. To study expression of *A. agallocha CYPs* in infected and healthy tree, raw RNA-seq data (Accession number PRJNA449813) in FASTQ format were obtained from NCBI SRA database. HISAT2-StringTie-DESeq2 workflow was utilized to calculate the differentially expressed members (Kim et al. 2015, Kovaka et al. 2019, Love et al. 2014). Library normalization was done based on median of ratios (Anders et al. 2010) with p ≤ 0.05. Differentially expressed genes (DEGs) were analysed for their pathway enrichment using ShinyGO 0.76.1 (Ge et al. 2020).

### 2.6. RNA extraction and cDNA conversion

Total RNA from wood tissue was extracted using modified CTAB method (Islam and Banu 2019), and from callus tissue with RNeasy Plant Mini Kit (Qiagen). Following DNase treatment, the integrity of extracted RNA was checked in 2% agarose gel and also estimated spectrophotometrically with Multiskan Sky Microplate Spectrophotometer (Thermo Fisher Scientific, US). SuperScript® III First-Strand Synthesis System (Invitrogen, USA) was used for cDNA preparation.

### 2.7. Real-time polymerase chain reaction (qRT-PCR) and statistical analysis

Real-time PCR was performed with Quant studio 3 Applied Biosystem, USA using SYBR green chemistry (Takara, Japan) to validate 10 candidate *AaCYP* genes for their differential expression in infected *Aquilaria* wood and MeJ induced callus. Primers were designed and optimised in Integrated DNA technology server (IDT) and synthesised. The reaction mix consisted of 0.5 μl (10pm/μl) each of the primer, 2 μl cDNA template, 10 μl SYBR Premix Ex Taq II (Takara, Japan), and nuclease-free water. The cycle parameters were set as 94 °C for 10 min during hold stage, 94 °C for 15 s and 58 °C for 60 s during PCR stage. Data normalization was done using Ct values of housekeeping gene (*GAPDH*) and relative expression was evaluated through 2 ^− ΔΔCt^ method. A oneway anova test with a 0.05 p-value was performed on three replicates (biological and technical) for each experiment.

### 2.8. Protein-protein interaction (PPI) and homology modelling

A network of interaction of differentially expressed AaCYPs were predicted based on proteinprotein connectivity network integrated in STRING v11.5 program with default parameters. 3D modelling was carried out with PHYRE2 server (Kelley et al. 2009). The models were investigated for quality assessment (Ray et al. 2012), pocket (Guilloux et al. 2011) and active site prediction (Wass et al. 2010).

## 3. Results

### 3.1. Identification of Cytochrome P450 family members in *Aquilaria agallocha*

Mining of *Aquilaria agallocha* genome resulted in the identification of 136 full-length CYPs, and their respective closest matched from UniProt database along with their deduced properties were elucidated (Additional file 1). The residue number of AaCYPs ranged lowest 338 (AaCYP69) to highest 821 (AaCYP40) and molecular weight from 43.66 KDa (AaCYP69) to 92.54 KDa (AaCYP40). The isoelectric point from 5.1 (AaCYP36) to 8.8 (AaCYP74), and transmembrane helices varied from zero in 56 AaCYPs, one in 66 AaCYPs, two in 11 AaCYPs, three in two AaCYPs, and four in one AaCYPs. Out of 136 AaCYPs, majority were located in endomembrane system (ES), seven in cytoplasm (C), five in organelle membrane (OM), and three each in plasma membrane (PM), chloroplast outer membrane (CM) and nucleus (N), and only one in mitochondrial membrane (MM).

### 3.2. Phylogenetic analysis and classification

Phylogenetic positions of the AaCYPs with CYPs of *Arabidopsis thaliana* (AtCYPs) enabled us to identify the close members based on sequence identity and classify them into clans and families (Fig. 1). The members were distributed in eight clans, one A-type i.e., CYP71 clan and seven non-A type i.e., CYP51, CYP72, CYP74, CYP85, CYP86, CYP97, and CYP711 clan. The four clans viz. CYP71, CYP85, CYP86, and CYP72 were observed to be multi-family clans with fifteen, nine, five, and four families, respectively. The CYP71 clan had the highest members with 82 AaCYPs (Additional file 2). The family members were further classified into subfamilies based on their position in the tree. Within CYP71 clan, CYP71 being the largest family had CYP71A subfamily (5 members), CYP71D subfamily (19 members), and CYP71E subfamily (1 member) (Fig 1). Interestingly, CYP83 members had clustered with CYP71 family. The CYP82 family had three subfamilies viz. CYP82A subfamily (10 members), CYP82D subfamily (one member), and CYP82G subfamily (one member). The CYP81 family was further divided into CYP81C subfamily (two members), CYP81E subfamily (two members), and CYP81Q (two members). The remaining families of CYP71 clan were not diversified into multiple subfamilies. Within CYP86 clan, three CYP704 family members belongs to CYP704C subfamily, two CYP94 family members distributed one each in CYP94A and CYP94C subfamily, and three CYP86 family members distributed each in CYP86A and CYP86B subfamily. Within CYP85 clan, the CYP716 family had two subfamilies viz. CYP716A (one member) and CYP716B (two members), CYP90 family had subfamilies viz. CYP90C (one member) and CYP90D (two member). Interestingly, the two CYP724 family members which formed cluster with CYP90 family, and were not named as CYP90 members in *A. thaliana* which indicates that CYP724 members in *A. agallocha* might had evolved from CYP90 family. The CYP97 clan members were distributed in three subfamilies viz. CYP97A, CYP97B, and CYP97C with one member each, and CYP74 clan had members in two subfamilies i.e., CYP74A (two members), and CYP74B (one member).

**Fig 1.**
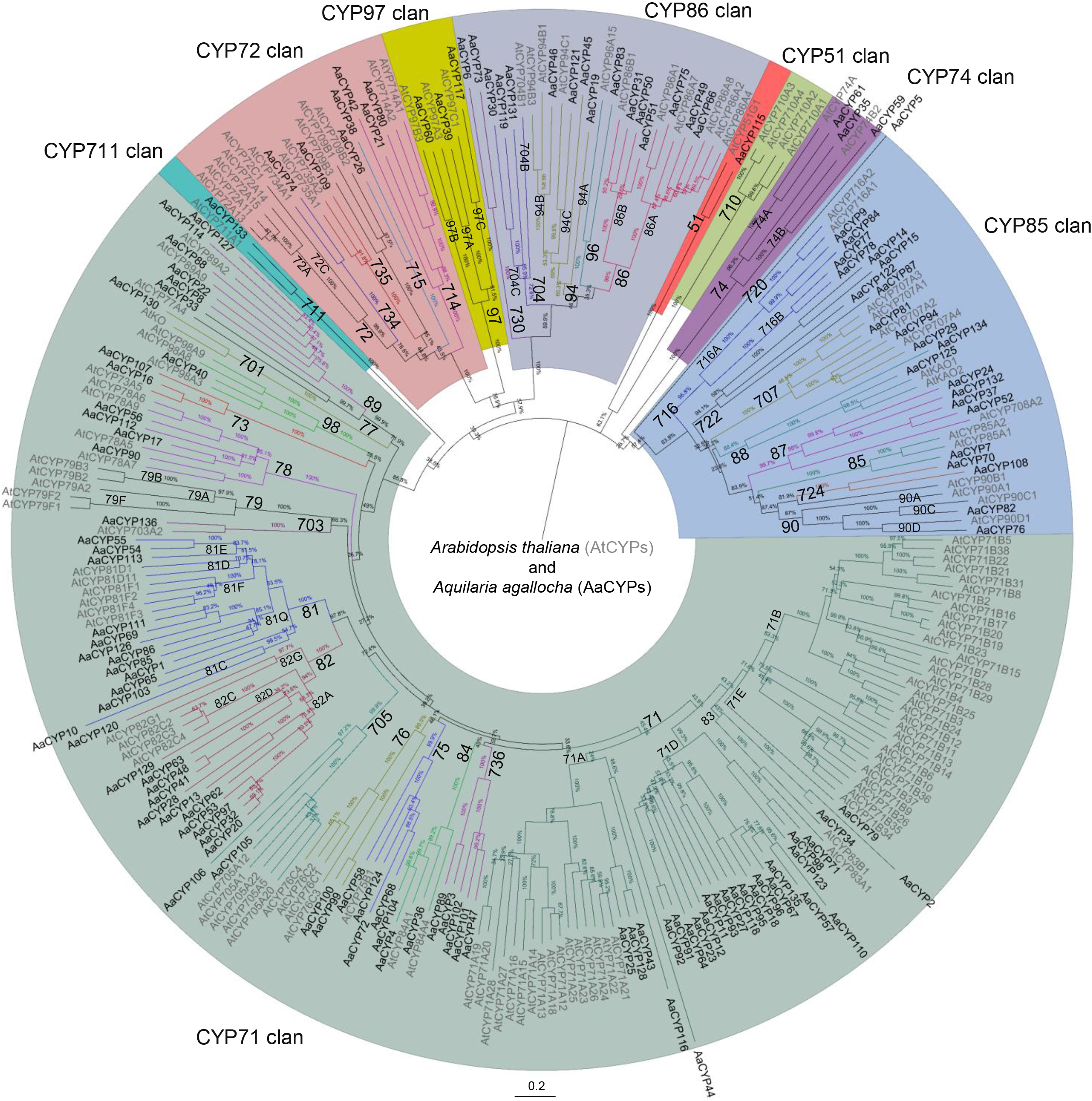
Phylogenetic positions of 136 Aquilaria agallocha CYPs with Arabidopsis thaliana CYPs (AtCYPs). The AaCYPs distributed into 8 clans were represented in different colour shades, and families within each clan were labelled beside the branch nodes.

### 3.3. Gene structures and their regulatory elements

The coding regions of *AaCYPs* were separated by a minimum of zero and maximum of sixteen introns (Fig. 2). The members of CYP97 clan had the highest number of introns (eleven to sixteen), followed by CYP85 clan (two to eight), and CYP72 clan (two to five). Few members of CYP71, CYP86, and CYP74 clan possessed no introns in their coding regions which shows that CYPs in *A. agallocha* vary greatly in terms numbers of intron-exon. The promoter sequence of the members had light-responsive, abiotic and biotic stress, hormone-responsive, site-binding, development-related, and unknown elements (Table 1 and Additional file 3). Presence of hormone and stress related elements suggest their inducible nature during stress and hormone exposure. However, 15.73% of CREs detected were unknown as no information are available in the Plant CARE database.

**Fig 2.**
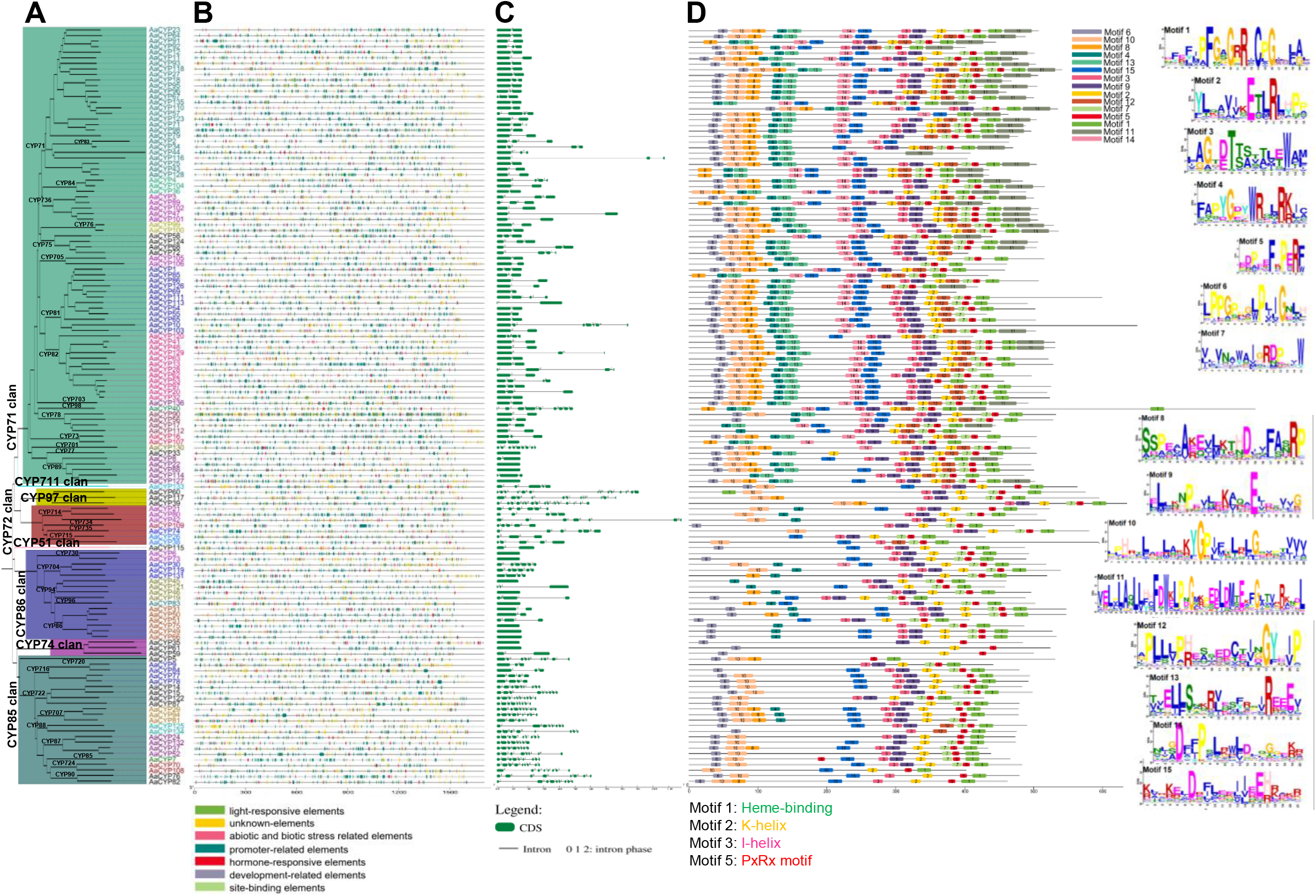
Presentation of phylogenetic tree, cis-regulatory elements (CREs), exon-intron boundaries, and conserve motif in the AaCYPs. **A)** Neighbor-joining tree constructed using protein sequences of AaCYPs. **B)** CREs in the 2000bp upstream regions of the AaCYPs. **C)** Gene structures (exon-intron boundaries). **D)** Conserved motifs in the AaCYP protein sequences. Motifs 1 to 15 with different colour shades were presented in amino acid level (right side), where larger residue size indicates more conserved.

**Table 1.** Categorization and types of Cis-regulatory elements (CREs) identified in the promoter regions of *AaCYPs*

| Sl. No | Element category | Number | Cis-regulatory elements |
| --- | --- | --- | --- |
| 1 | Promoter related | 9095 | TATA-box and CAAT-box |
| 2 | Unknown | 2410 | - |
| 3 | Light-responsive | 1392 | GATA-motif, TCT-motif, GT1-motif, Box 4, G-box, ACE, I-box, TCCC-motif, MRE, LAMP-element, Sp1 |
| 4 | Abiotic and biotic stress | 1189 | WUN-motif, WRE3, as-1, STRE, TC-rich repeats, GC-motif, box S, DRE1, ARE, DRE core, MBS, LTR, STRE |
| 5 | Hormone-responsive | 1017 | TCA-element, ERE, TCA, CGTCA-motif, TGACG-motif, AT~ABRE, ABRE, ABRE3a, ABRE4, GARE-motif, P-box, AuxRR-core, TGA-element |
| 6 | Development-related | 202 | CAT-box, O2-site, GCN4_motif, CCGTCC-box, motif I, circadian, RY-element, AACA_motif, ref2f-1, HD-Zip1, AT-rich element, MSA-like |
| 7 | Site-binding | 16 | CCAAT-box and E2Fb |

### 3.4. Motif composition and conserved residues

Motif analysis in the AaCYPs revealed heme-binding (motif 1, 92.64% AaCYPs); K-helix (motif 2, 88.97% AaCYPs); I-helix (motif 3, 91% AaCYPs); and PXRX (motif 5; 90% AaCYPs) to be conserved (Fig. 2). Additionally, 11 other motifs were also detected in 52.2% to 91.9% of AaCYPs (Fig. 2). However, the residues in the conserved regions tend to vary between A-type and non-A type CYPs. For e.g., in A-type clan, consensus residues of the heme-binding motif were PFGXGRRXCPG, while in non-A type clans, F/YXXFXXGXRXCXG were consensus (Fig 3). Similarly, the consensus residues of K-helix were ET/SLR in A-type members, while EXLR in non-A type members. The PXRX motif appeared consensus as FXPERF in A-type and FXPXRF/W in non-A type clan members. Similarly, consensus residues of I-helix were A/GGXD/ETS/T in A-type and AGXD/ET/S in non-A type. The helix was the most conserved, however missing in CYP74 clan members which probably because of their NADPH and molecular oxygen independent catalytic nature. The members of same subgroups showed similar motif composition, but varies between family members which indicates their association with diverse functions.

**Fig 3.**
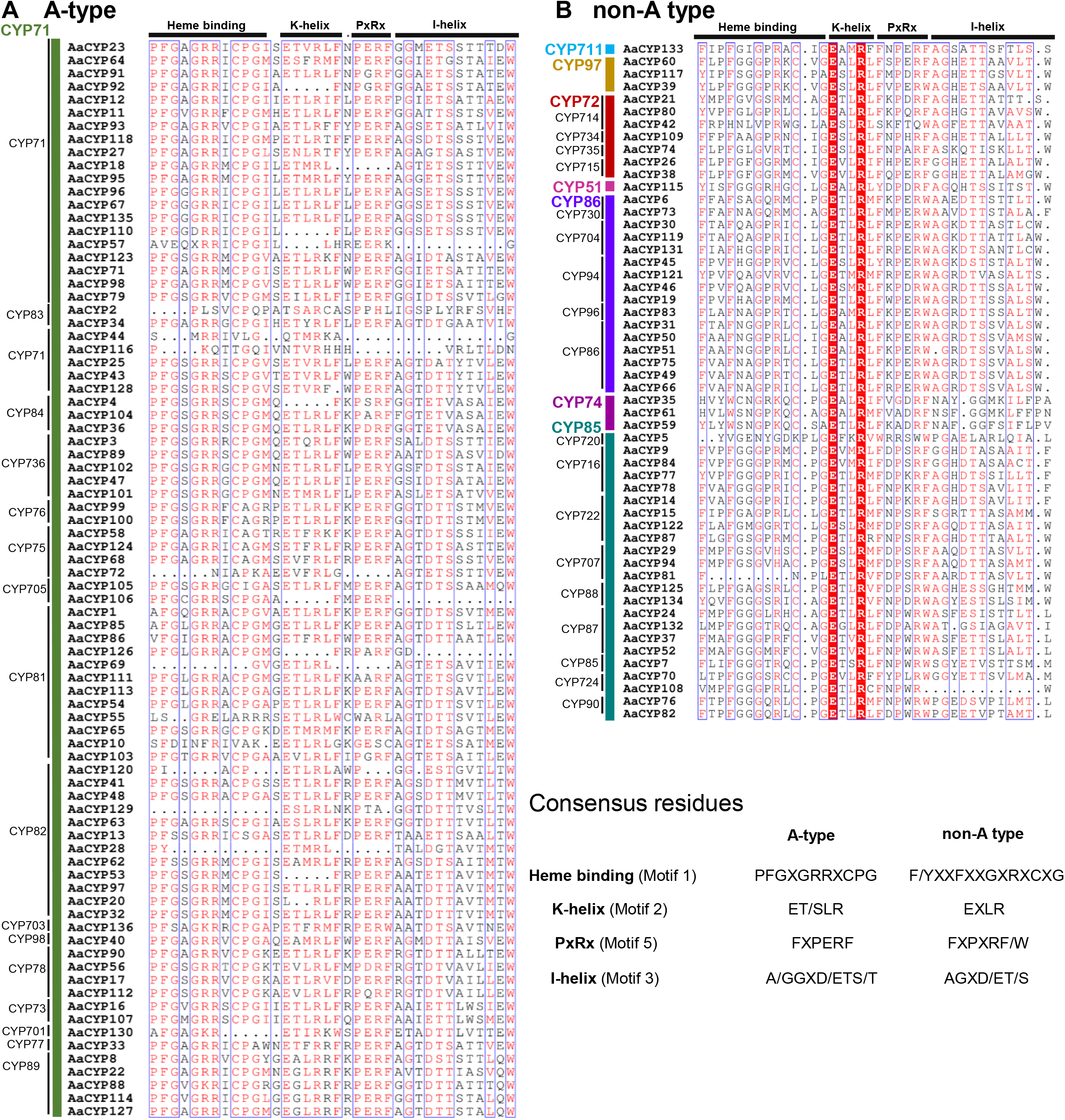
Presentation of the family-wise amino acid residues of the four most conserved regions (Heme-binding, K-helix, PxRx, and I-helix) in **A)** A-type and **B)** non-A type.

### 3.5. Synteny and duplication analysis

The *AaCYPs* in the genome were distributed in 119 scaffolds. Collinearity analysis led to the identification of evolutionary related gene pairs with *V. vinifera, A. thaliana, S. lycopersicum*, and *G. max* (Fig. 4). Highest number of orthologous genes matched with *G. max* (53 pairs), followed by *V. vinifera* (49 pairs), *S. lycopersicum* (29 gene pairs), and *A. thaliana* (16 pairs) (Additional file 4). In *G. max*, the members were dispersed in 18 of 20 chromosomes where most pairs were associated between chromosome 11 and scaffold KK899083.1 of *A. agallocha* (Fig 4). Gene pairs of *A. thaliana* were present in all the chromosomes, with highest number of genes located in chromosome 1 and 2 and scaffold KK898996.1 of *A. agallocha*. In *S. lycopersicum*, orthologous pairs with *A. agallocha* were located in chromosome 1-4 and 7-10, chromosome 2 hold maximum number of pairs with a total of nine scaffolds. In *V. vinifera* orthologous pairs were dispersed in 18 of 19 chromosomes. Scaffold KK898996.1 showed highest number of pairs which were dispersed across chromosome 1, 2, 3 and 15 of *V. vinifera*. Interestingly, nine AaCYPs which belongs to CYP71, CYP72, and CYP85 had orthologous pair with all the four plants (Fig. 4), where *AaCYP78A1* possessed highest in number indicating that it might have a crucial role in metabolism. Duplication analysis led to identification of 10 pairs of segmental (SD) and 4 pairs of tandem duplicated genes (Additional file 5). The Ka/Ks ratio ranged from 0.080 (*AaCYP86A1/AaCYP86A2*) to 0.345 (*AaCYP71D10/AaCYP71D11*), while the MYA ranged from 13.76 (*AaCYP82A7/AaCYP82A10*) to 140.70 (*AaCYP86A1/AaCYP86A2*) (Additional file 6).

**Fig 4.**
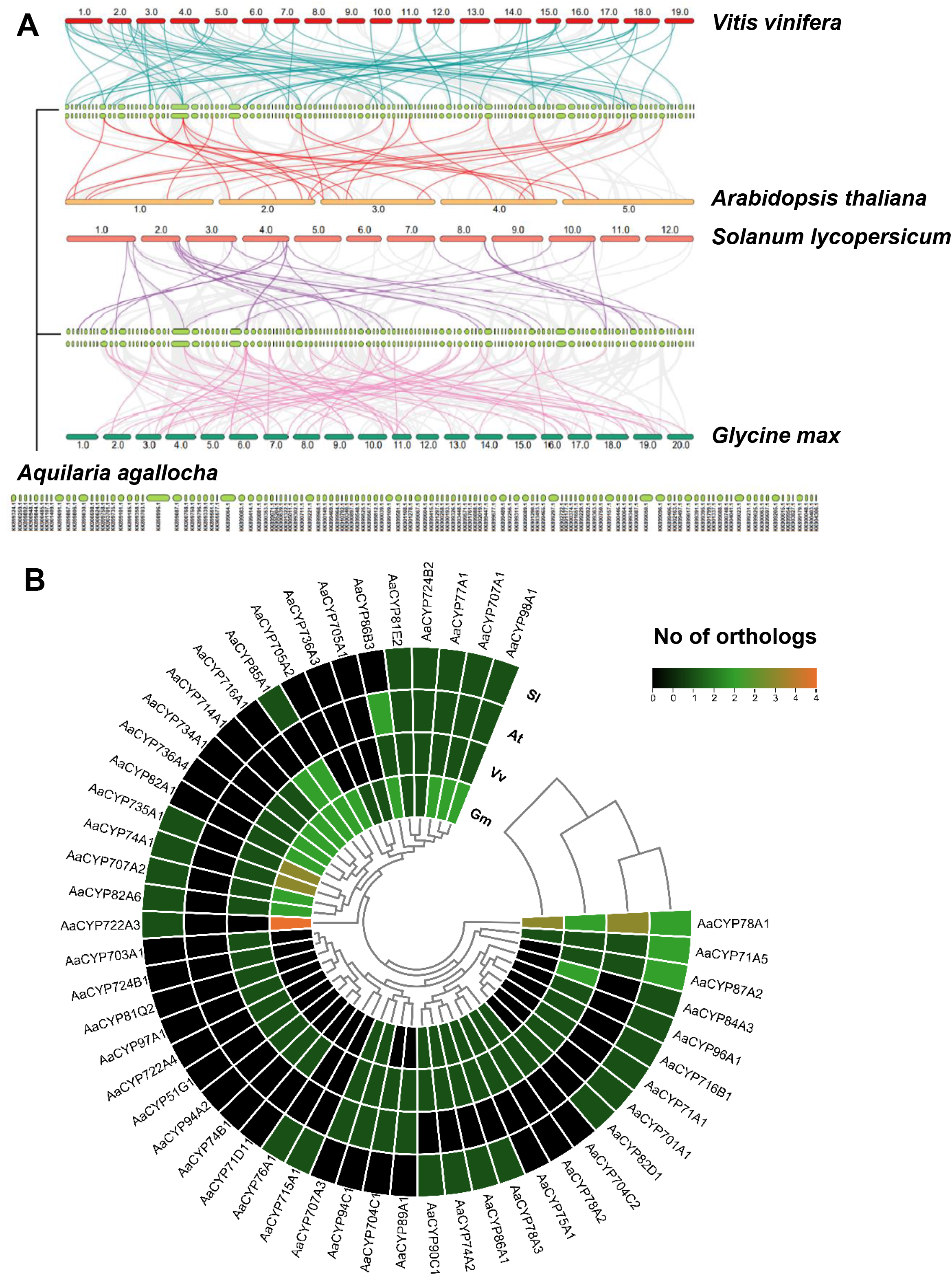
Overview of evolutionary relationship of CYPs of A. agallocha with V. vinifera, A. thaliana, S. lycopersicum, and G. max. **A)** Synteny relation between AaCYPs and four other angiosperms. The grey lines between the genomes represents the collinear blocks, and the different coloured lines represents syntenic CYPpairs. **B)** Number of ortholog of AaCYPs (syntenic pairs) with CYPs of *V.vinifera* (Vv), *A. thaliana* (At), *S. lycopersicum* (Sl), and *G.max* (Gm).

### 3.6. Functional categorization and expression analysis

Gene Ontology (GO) assignment of AaCYPs revealed their association in various biological process (BP), cellular components (CC) and molecular functions (MF) (Fig. 5). The percentages of AaCYPs associated with biological processes were: multicellular organismal process (6.9%), development process (6.5%), anatomical structure was development (6.5%), response to stress (5.8%), hormone metabolic process (5.1%), response to biotic stimulus (4.7%), and so on (Fig. 5 and Additional file 7). As expected, in molecular function, 92.9% showed oxidoreductase activity, 2.4% showed lyase activity, and 1.2% each of oxygen binding, transferase and demethylase activity. Out of 136 AaCYP proteins, only 46.6% could be mapped into the pathways defined in KEGG (Additional file 8) where 14% involved in terpenoids and polyketides (diterpenoid, carotenoid, brassinosteroid, and zeatin) biosynthesis; 14% in other metabolites (phenylpropanoid, flavonoid, flavone, flavonol, and isoflavonoid, stilbenoid, diarylheptanoid) biosynthesis; 11% in lipid (fatty acid, cutin, suberin, wax, steroid, and alpha-linolenic) metabolism; and only 2% in cofactors and vitamins (ubiquinone, terpenoid-quinone) metabolism.

**Fig 5.**
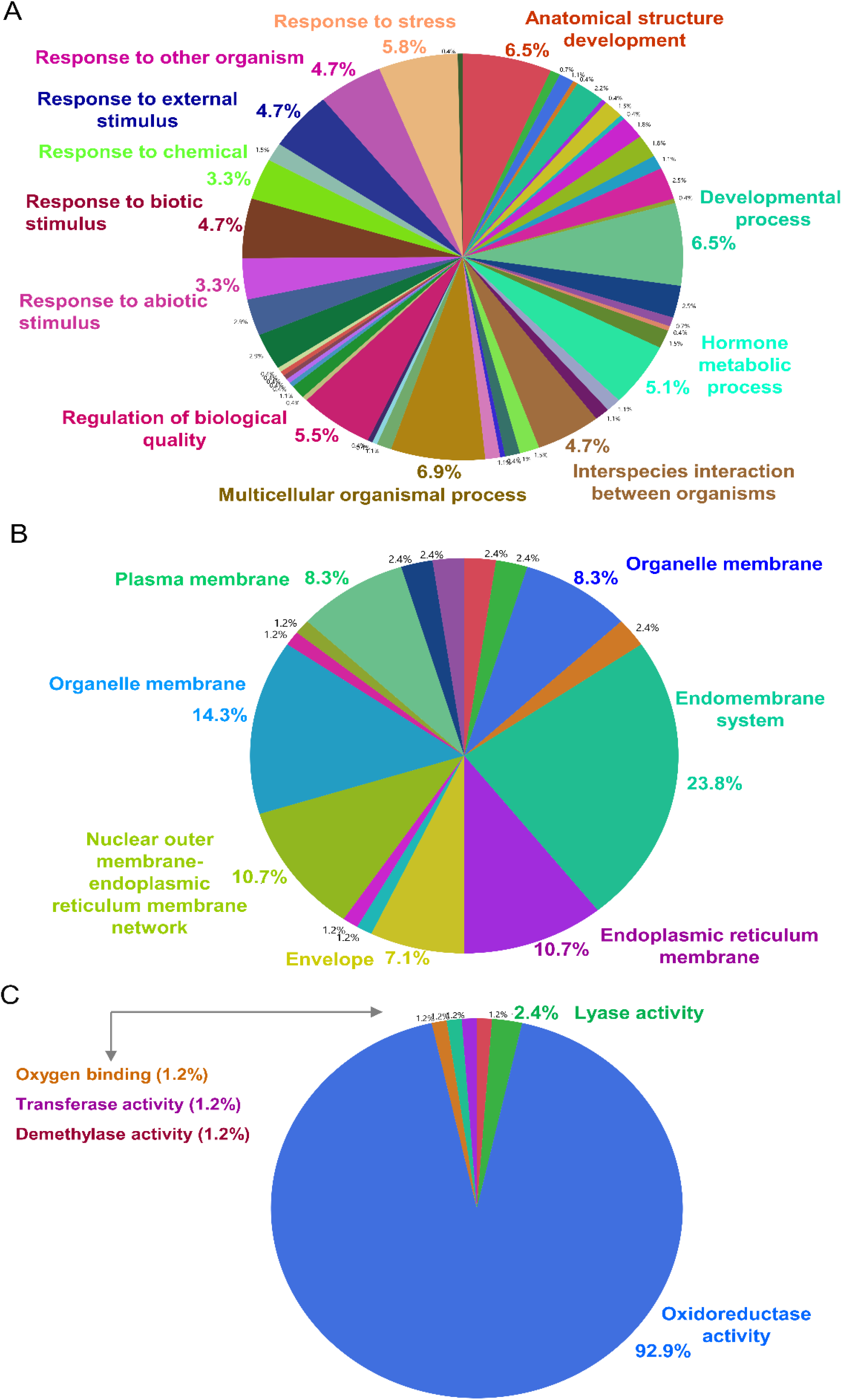
Functional prediction and categorization of 136 AaCYPs. Percentage of AaCYPs involved in various **A)** Biological processes. **B)** Cellular components. **C)** Molecular functions.

RNA-seq data analysis revealed 76 *AaCYPs* to be differentially expressed (DEGs) in stress induced *Aquilaria* when compared with normal healthy tree (Fig. 6). The DEGs belongs to clans CYP71 (50 members), CYP72 (5 members), CYP85 (12 members), CYP86 (6 members), and CYP97 (1 member) (Additional file 9). Pathway mapping revealed the functional positions of these DEGs. For instance, through phenylpropanoid pathway, simple phenolics such as ferulic acid, sinapic acid, coniferylalcohol, and syringin reported from *Aquilaria* were biosynthesized, where the members AaCYP73A1 (2.4 log2FC), and AaCYP73A2 (5.78 log2FC) probably catalyse trans-cinnamate and cinnamoyl-CoA into p-coumaric acid and p-coumaroyl-CoA. (Fig. 7). The member AaCYP98A1 with log2FC of 1.24 was responsible for conversion of 4-coumaroylshikimate into caffeoyl shikimic or chlorogenic acid. Two CYP84 family members viz. AaCYP84A1 (6.99 log2FC) and AaCYP84A2 (4.24 log2FC) probably convert simple phenolics such as ferulic acid to 5-hydroxyferulic, coniferylaldehyde to 5-hydroxyconiferal aldehyde, and coniferylalcohol to 5-hydroxyconiferal alcohol (Fig. 7). Similarly, the members of CYP75 family were involved in generation of metabolites including kaempferol, butin, fustin, quercetin, apigenin, sakuranetin, luteolin, and myricetin (Fig.7), where AaCYP75B1 (7.90 log2FC) and AaCYP75B3 (6.21 log2FC) probably accept substrates such as liquirtugenin, garbanzol, apigenin, naringenin, and convert them into flavonoid derivatives (Fig. 7). Three members AaCYP724B1, AaCYP734A1, AaCYP90D1 of brassinosteroid biosynthesis were upregulated by 6.12, 5.56, and 3.97 log2FC, respectively (Fig. 6). Additionally, members in biosynthesis of carotenoid (AaCYP707A1), isoflavonoid (AaCYP81Q), stilbenoid, diarylheptanoid and gingerol (AaCYP73A2, AaCYP98A1), fatty acid degradation and wax biosynthesis (AaCYP86A3), alpha-linolenic metabolism (AaCYP74A1) were elevated 1.24 to 9.74 log2FC (Fig. 6). However, a member each from diterpenoid biosynthesis (AaCYP701A1), and steroid biosynthesis (AaCYP51G1) and two from carotenoid (AaCYP707A2, AaCYP707A3) were downregulated by 1.09, 3.62, 1.60, and 3.33 log2FC, respectively (Fig. 6).

**Fig 6.**
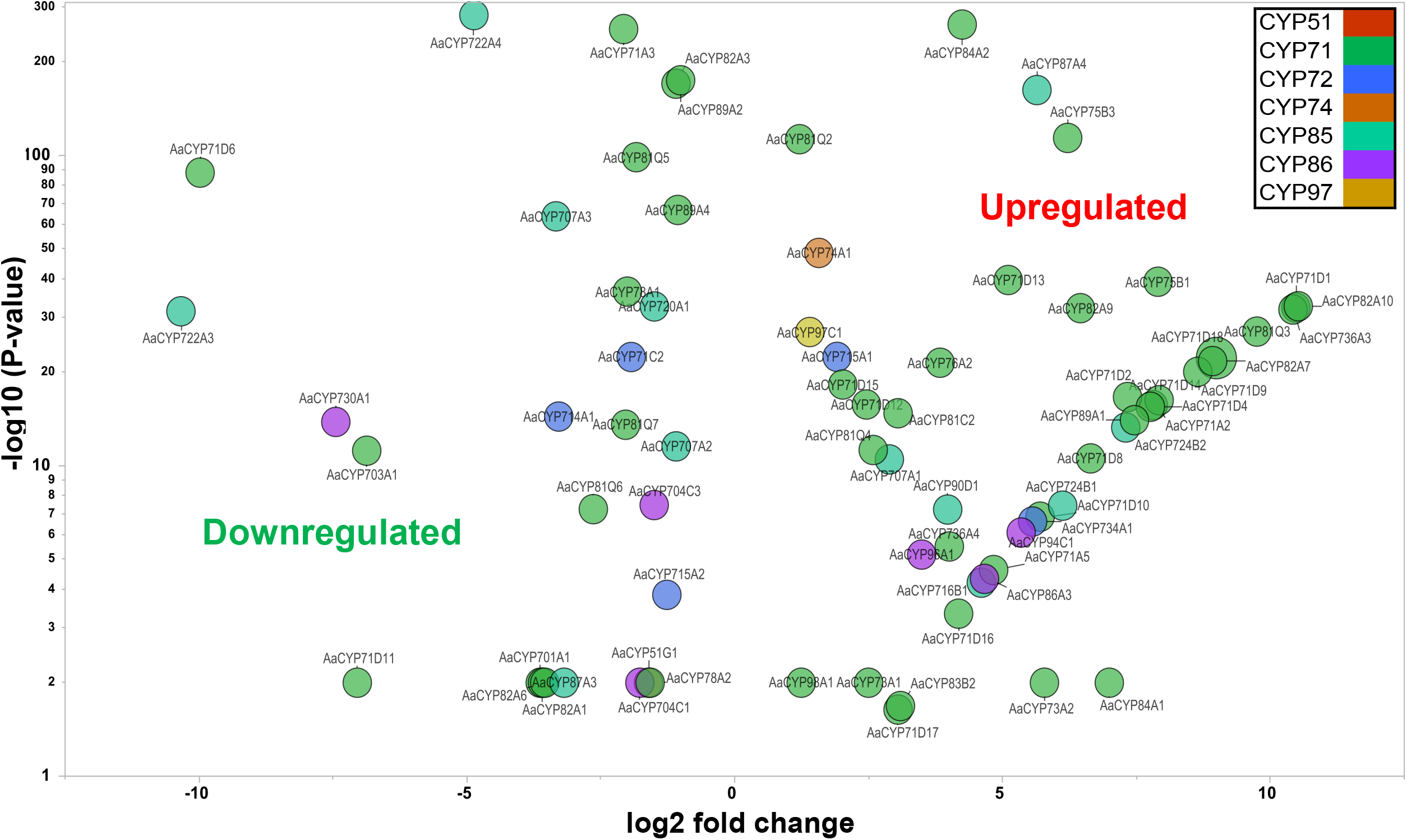
Differentially expressed AaCYPs in stress induced A. agallocha compared to healthy tree. X and Y-axis represents the log2 fold change (log2FC) and negative log p-value respectively. The clan members were presented in different colour shades.

**Fig 7.**
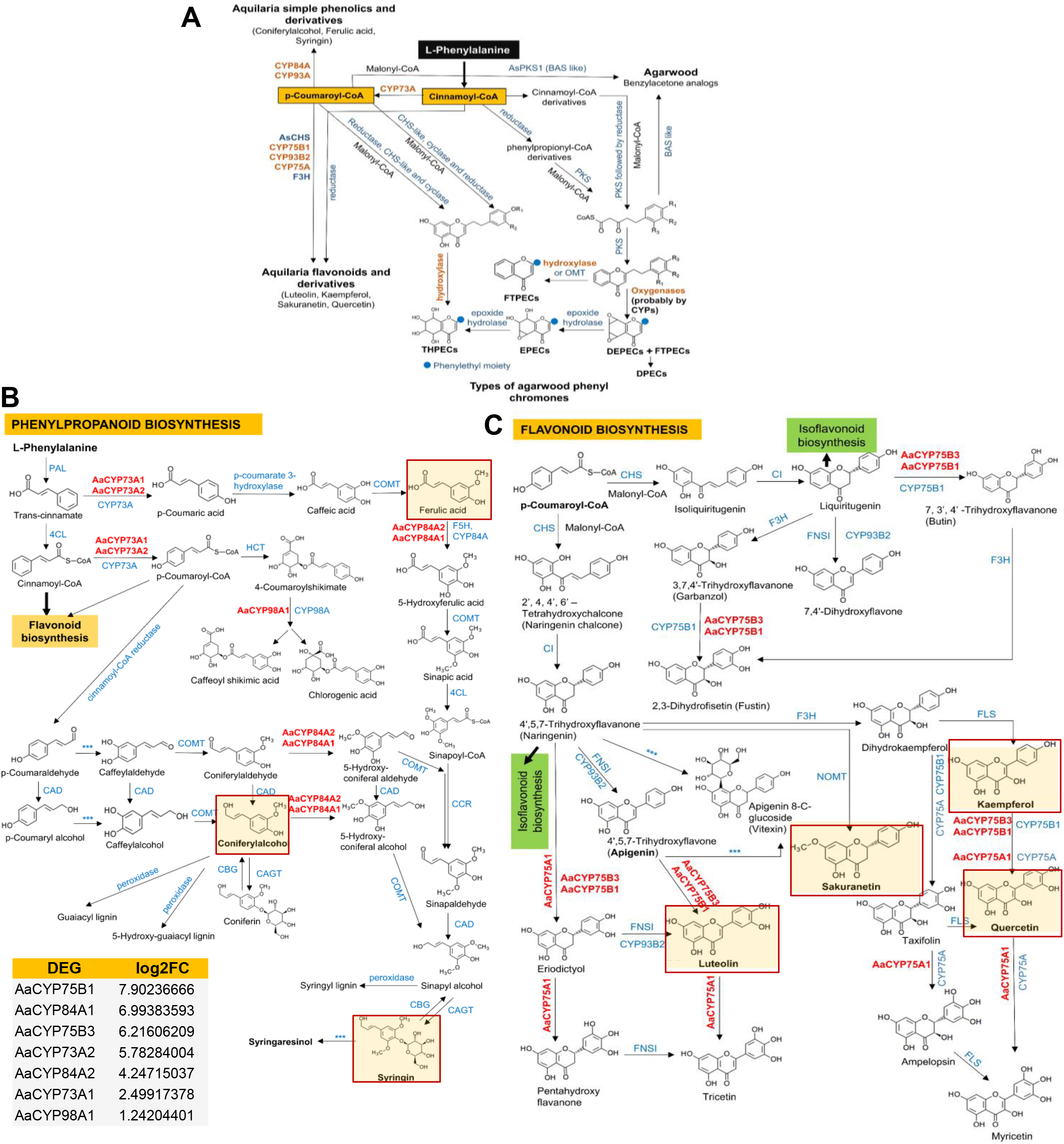
Proposed pathway and CYPs involvement in biosynthesis of secondary metabolites in A. agallocha. **A)** Involvement of CYP73A, CYP75A, CYP75B1, CYP84A, CYP93A, and CYP93B2 for phenolics, flavonoids and phenylethyl chromones biosynthesis from the common precursors (p-Coumaroyl-CoA and Cinnamoyl-CoA). Partners in the pathway are chalcone synthase (CHS), benzalacetone synthase (BAS), polyketide synthase (PKS), flavonoid 3□ hydroxylase (F3H), reductases, and cyclases. The hydroxylases (probably by CYPs) and oxygenases (probably by CYPs), O-methyltransferase (OMT) and epoxide hydrolases catalyses the conversion between the type of phenylethyl chromones i.e., Flindersia type (FTPECs), diepoxy-5,6,7,8-tetrahydro type (DEPECs), mono-epoxy-5,6,7,8-tetrahydro type (EPECs), 5,6,7,8-tetrahydro type (THPECs), dimeric type (DPECs), trimeric type (TPECs) identified in *Aquilaria* (Chen et al. 2012 and Li et al. 2021). **B)** Biosynthesis of phenolics and derivatives by AaCYP73A1, AaCYP73A2, AaCYP84A1, AaCYP84A2, and AaCYP98A1 through phenylpropanoid pathway. The compounds identified in *Aquilaria* plants were presented in red colour box, and AaCYPs involved in the biosynthesis are presented in red colour text. **C)** Biosynthesis of flavonoids such as luteolin, kaempferol, sakuranetin, and quercetin (marked red box) by AaCYP75A1, AaCYP75B1, and AaCYP75B3. The un-marked compounds have not yet identified in *Aquilaria* and *** indicates enzymes characterization not done.

Since only 23% of the DEGs could be mapped to KEGG, review of information from UniprotKB (sequence similarity) and literature enabled us to predict the functions of the remaining DEGs (Additional file 9). Interestingly, AaCYP82A10 and AaCYP82A7 which were upregulated by 9 to 10 log2FC associated with pathogen resistance, biotic and abiotic stress (Fig. 6). AaCYP71D7 and AaCYP71D17 were involved in biosynthesis of sesquiterpenoid (Fig. 8) and the later was upregulated by log2FC of 3.05. Two additional members AaCYP724B2 and AaCYP715A2, involved in brassinosteroid biosynthesis and brassinosteroid inactivation upregulated (log2FC 7.3) and downregulated (log2FC 1.26) respectively (Fig. 6). However, members involved in processes such as development of shoot apical meristem development (AaCYP78A2); gibberellin deactivation (AaCYP71C2, AaCYP714A1); abscisic acid degradation (AaCYP722A3, AaCYP722A4) were downregulated significantly. Moreover, AaCYP86A3 which was involved in biosynthesis of cell wall compounds (cutin, suberin, wax) and oxidation of fatty acid for degradation, upregulated by log2FC of 4.66, while AaCYP703A1, probably the hydroxylase for fatty acid metabolism downregulated by 6.86 log2FC. The expression values and pathways of all the members were summarized in (Additional file 9).

**Fig 8.**
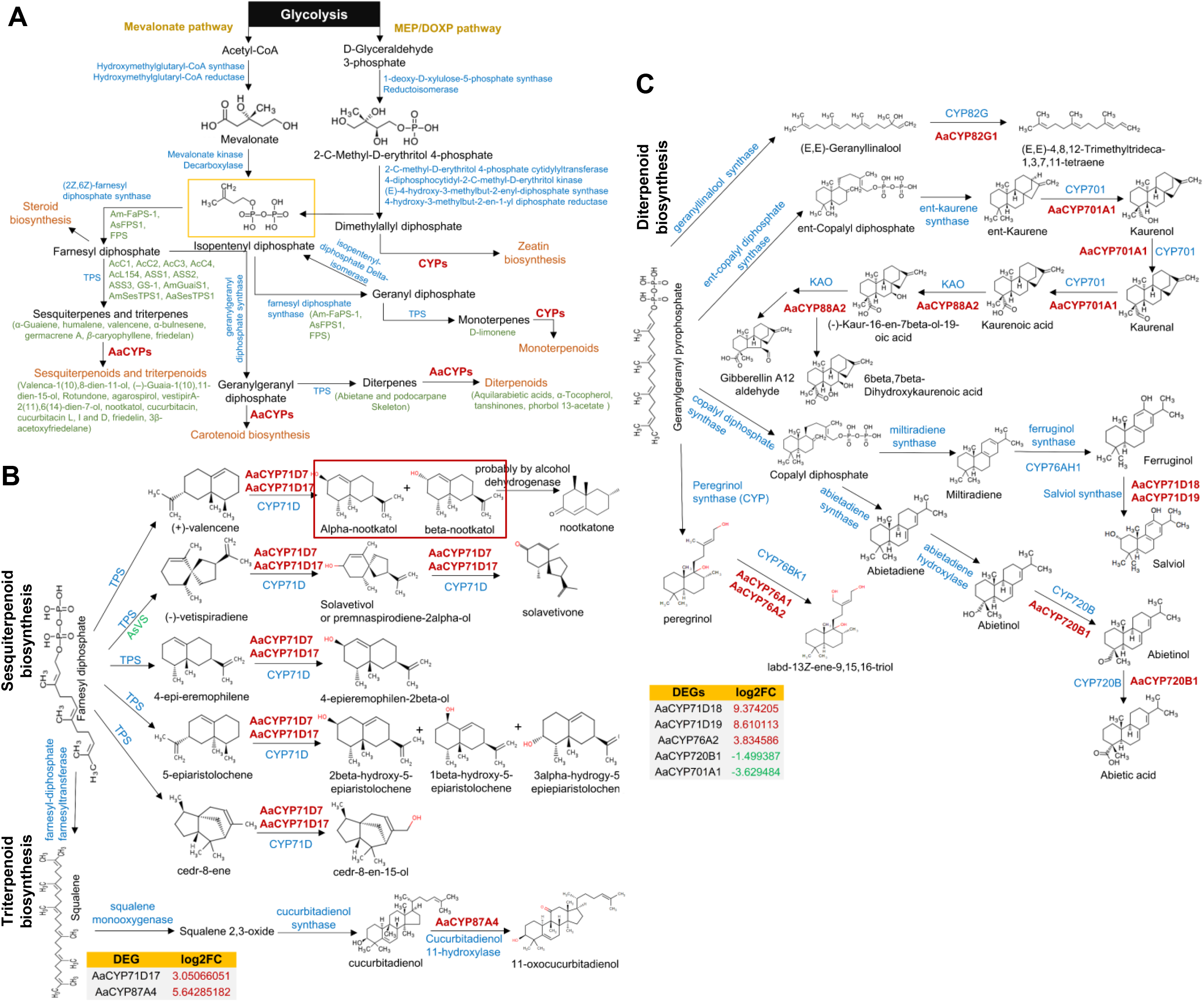
Overview of CYPs involvement in the biosynthesis of terpenoids and other metabolites in A. agallocha. **A)** The common pathway for the biosynthesis of terpenoids, steroids, zeatin, and carotenoids. The olive-green colour text indicates the proteins and metabolites identified in different species of Aquilaria and TPS and FPS stands for terpene synthases and farnesyl diphosphate synthase. **B)** AaCYP71D7 and AaCYP71D17 of A. agallocha probably act as hydroxylase and oxygenase in biosynthesis of nootkatol (identified in agarwood) and other sesquiterpenoids. **C)** AaCYPs involvement in diterpenoid biosynthesis, where AaCYP720B1 involved in abietic acid (abietane have been detected in agarwood) biosynthesis. Other members in the pathway are CYP71D, CYP82G, CYP88A, and CYP701A possibly involved in salviol and peregrinol (so far not identified in agarwood) biosynthesis. **D)** The member AaCYP87A4 mapped into triterpenoid biosynthesis which probably has hydroxylase activity towards triterpenoids.

### 3.7. Pathway enrichment analysis and interaction network

Pathway enrichment analysis helped identify the pathways which were enriched in the DEGs (Table 2). The upregulated DEGs which belongs to flavone and flavonol, flavonoid, and brassinosteroid biosynthesis were enriched with fold enrichment of 158.1, 79.1 and 26.4 respectively. Whereas, downregulated DEGs of diterpenoid and steroid pathway enriched by 60.2 and 35.1-fold respectively. Interestingly, both upregulated and downregulated DEGs were involved in enriched pathways such as carotenoid, cutin, suberin and wax, and steroid biosynthesis.

**Table 2.** Enriched pathways of Differentially expressed genes (DEGs) in induced *A. agallocha*

| DEGs regulation | Enrichment FDR | Fold Enrichment | Pathways |
| --- | --- | --- | --- |
| Up | 1.3E-02 | 158.1 | Flavone and flavonol biosynthesis |
|  | 2.0E-02 | 79.1 | Brassinosteroid biosynthesis |
|  | 3.5E-03 | 43.6 | Carotenoid biosynthesis |
|  | 3.8E-03 | 34.2 | Cutin, suberin and wax biosynthesis |
|  | 5.0E-02 | 26.4 | Flavonoid biosynthesis |
|  | 2.7E-03 | 3.6 | Biosynthesis of secondary metabolites |
| Down | 2.9E-02 | 60.2 | Diterpenoid biosynthesis |
|  | 2.9E-02 | 43.6 | Carotenoid biosynthesis |
|  | 2.9E-02 | 35.1 | Steroid biosynthesis |
|  | 2.9E-02 | 34.2 | Cutin, suberin and wax biosynthesis |
|  | 1.8E-02 | 4.1 | Biosynthesis of secondary metabolites |

The network model consisted of 80 nodes and 364 edges with protein-protein enrichment significance of < 1.0e-16 (p-value). The interaction map had 4 distinct clusters derived from the protein’s involvement in various metabolic pathways (Fig 9). The members viz. AaCYP84A2, AaCYP98A1, AaCYP73A1, AaCYP75B3 and their partners in flavonoid and phenylpropanoid biosynthesis appeared as single cluster. Interestingly, AaCYP73A1, AaCYP98A1 and HCT of *A. thaliana* showed association with all the three i.e., flavonoid, phenylpropanoid, and stilbenoid pathways. The member AaCYP74A1 which appeared ortholog to allene oxide synthase (AOS) clustered with allene oxide cyclase (AOC1, AOC2) and lipoxygenase 2 (LOX2) (Fig 9), which probably contributes to the biosynthesis of methyl jasmonate. The third cluster composite of AaCYP51G1 participated in sterol biosynthesis with the partners in the pathway. The fourth cluster involves four members of brassinosteroid biosynthesis namely AaCYP90D1, AaCYP724B1, AaCYP724B2, and AaCYP734A1. Interestingly, the interaction that involved AaCYP94C1 (jasmonate-mediated signaling), AaCYP74A1 (jasmonate biosynthesis), PAL1 (phenylpropanoid biosynthesis) and TTR (flavonoid biosynthesis) suggest jasmonate mediated regulation of phenolics and flavonoids biosynthesis during stress in *Aquilaria* plant species.

**Fig 9.**
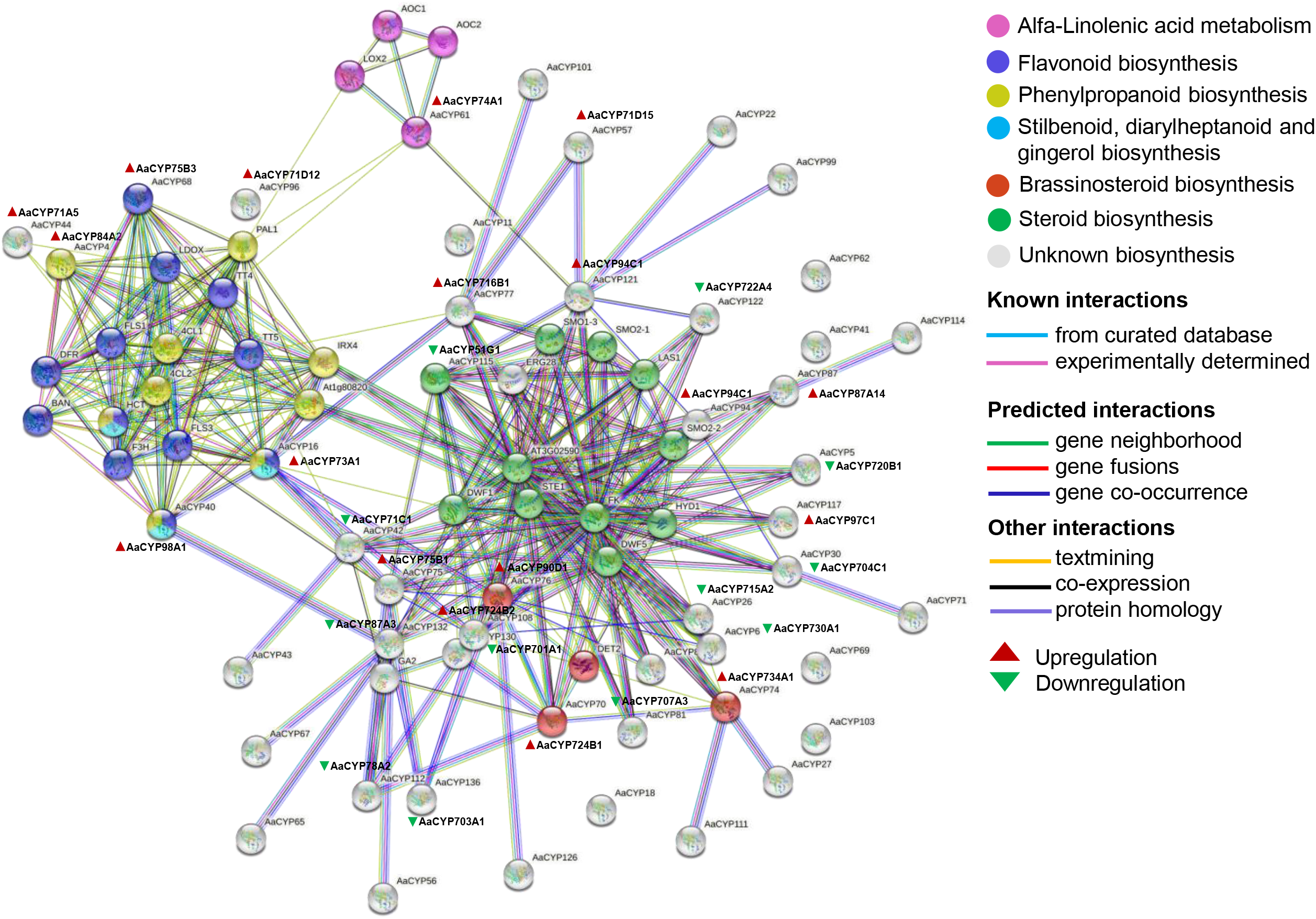
Network model of differentially expressed AaCYPs, where the potential AaCYPs with their functional partners in each enriched pathways are presented in similar colour, and their type of interactions are shown in different colour lines. The functional partners of AaCYPs are 1) In phenylpropanoid: ammonia-lyase 1, PAL1. 2) In flavonoid biosynthesis: chalcone synthase, TT4; chalcone-flavanone isomerase, flavanone 3-hydroxylase, FLS1; flavonol synthase 3, FLS3; dihydroflavonol reductase, DFR; 2-oxoglutarate 3-dioxygenase, F3H. 3) In steroid biosynthesis: sterol 4-alpha methyl oxidase (SMO1/2), lanosterol synthase 1 (LAS1), fatty acid hydroxylase (AT3G02590), cell elongation protein (DWF1), and hydroxysteroid isomerase (HYD1).

### 3.8. Validation of candidate *AaCYPs* in infected wood and methyl jasmonate (MeJ) treated callus

Selection of the candidate members were done considering their predicted involvement in phenylpropanoid biosynthesis (*AaCYP73A2, AaCYP84A1, AaCYP84A2*, and *AaCYP98A1*, and terpenoid biosynthesis (*AaCYP71D1, AaCYP71D4, AaCYP71D7, AaCYP71D10, AaCYP71D11*, and *AaCYP71D17*) (Additional file 10). In naturally infected tissue, all the 10 members were differentially expressed and upregulated significantly in the range of 6 to 13 log2FC compared to the healthy tissue (Fig. 10) indicating their role in phenolics and terpenoids biosynthesis during stress and possibly during agarwood formation. However, in MeJ treated callus tissue, *AaCYP71D7, AaCYP71D10*, and *AaCYP71D17* showed no changes in the expression. But rest of the 7 genes upregulated significantly in the range of 6 to 13 log2FC indicating MeJ role as an inducer in the biosynthesis of these metabolites.

**Fig 10.**
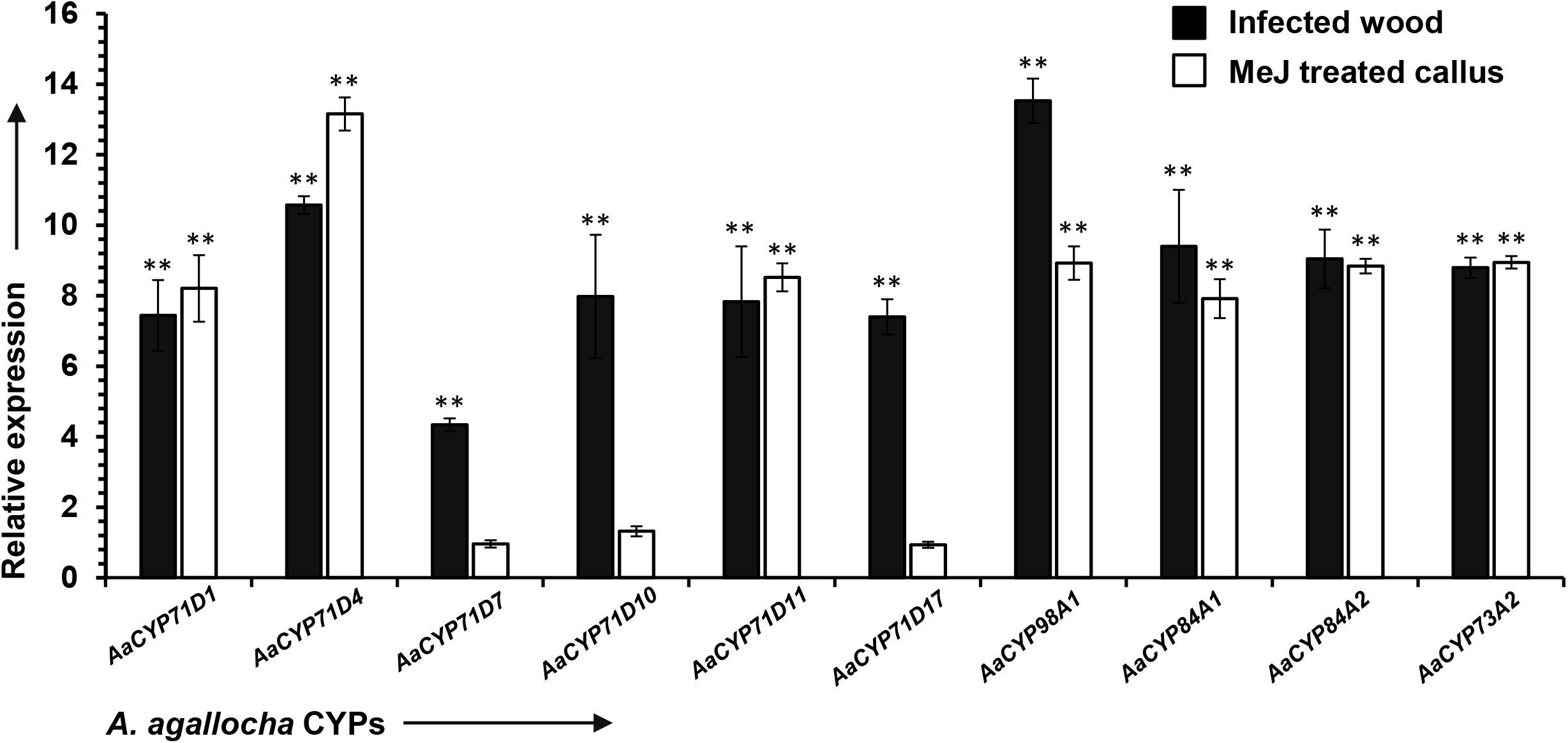
Expression profiles of 10 candidate *AaCYPs* which involved in sesquiterpenoid biosynthesis (*AaCYP71D1, AaCYP71D4, AaCYP71D7, AaCYP71D10, AaCYP71D11*, and *AaCYP71D17*), and phenylpropanoid biosynthesis (*AaCYP73A2, AaCYP84A1, AaCYP84A2*, and *AaCYP98A1*). Relative expression in infected A. agallocha tree (represented as infected wood) compared with healthy tree (non-infected wood), and in methyl jasmonate (MeJ) treated callus (represented as MeJ treated callus) compared with control. Statistical significance of the samples was obtained through a one-way anova, where ** indicates p-value < 0.01 and * indicates p-value < 0.05.

### 3.9. 3D modelling and active site prediction

Tertiary structures of AaCYPs were modelled with 100% confidence and 66 to 93% coverage (Fig 11). The contribution of secondary structures ranged from α-helix, 47-55%; β-strand, 8-11%; and TM helix, 2-13% including turns and loops. The structure of AaCYP51G1 and AaCYP724B1 had the maximum number of alpha helix (24) and beta strand (14) respectively. The root means square deviation (RMSD) between 0.085 to 0.013 Å indicates good quality of the models. Additionally, structure assessment analysis further verified the model quality and reliability (Additional file 11). The catalytic heme binding domain, appeared as loop; PXRX, appeared as loop in AaCYP97A1 and AaCYP74A1 but as helix in others (Additional file 12). The active amino acids associated in polar interactions were observed indicating their importance in the active pocket and in the conserved regions (Fig 11). Notably many non-polar and a few covalent interactions were also detected which probably stabilizes the complex during catalysis.

**Fig 11.**
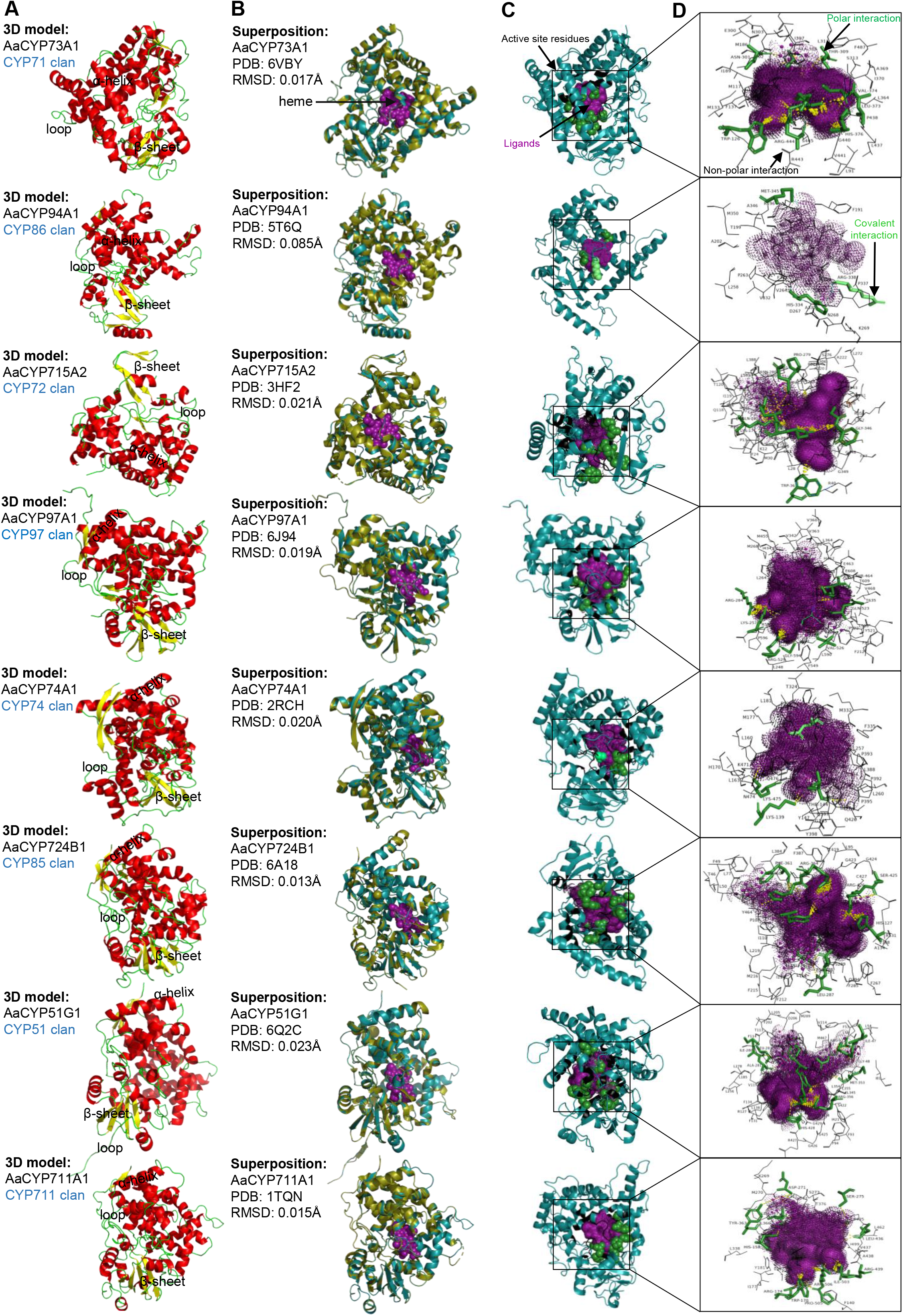
3D models of 8 AaCYP members from respective CYP clans. A) The 3D structures composed of α-helixes (indicated in red), β-sheets (indicated in yellow), and loops (indicated in green). B) 3D alignment of AaCYPs (deep teal) with their template PDB (olive green). C) Ligand or substrate-binding sites were predicted using server 3DLigandSite and indicated in black colour box. D) Structures visualized with PyMol, where residues involved in polar interaction (green stick), nonpolar interaction (black wire), and covalent interaction (pale green) were showed. The polar interaction of the predicted ligands with the AaCYP protein are depicted in yellow dashes.

## 4. Discussion

### 4.1. Evolution of AaCYPs and relation with other plants CYP members

In the genomes of Viridiplantae, CYPomes ranges from 0.2% to 1% of the protein-coding regions. Their diversity and functional role have been specifically described in plants (Hansen et al., 2021; Nelson and Werck-Reichhart, 2011) and their number starting from the simplest algae *C. reinhardtii* (39 CYPs), and moss *P. patens* (71 CYPs) to angiosperms including *A. thaliana* (246 CYPs), *O. sativa* (329 CYPs), *V. vinifera* (457 CYPs), *E. grandis* (443 CYPs), and *G. max* (332 CYPs) (Nelson and Werck-Reichhart, 2011; Nelson et al. 2009) suggest their diversification quite early during evolution. In the current study, 136 full-length CYP genes were detected in the 726 Mb genome of *A. agallocha* which were distributed into 8 clans and 38 families based on AtCYPs as reference. The total CYPs in *A. agallocha* were lowest among all the mentioned angiosperms while the genome was largest which supports the statement illustrated by Sun et al. 2020 that no correlation exists between the CYPs number and genome sizes. The AaCYPs which belongs to CYP71, CYP81 and CYP82 family were abundant i.e., 27, 12, and 12 in number respectively within A-type clan while, in non-A type clan, members of CYP86, CYP87, CYP716, CYP722 family were abundance. CYPs of *A. thaliana* were distributed in 47 families (Nelson et al. 2009), where CYP71, CYP81 and CYP705 family members were abundant while possess only five members of CYP82 family. In non-A type clans, CYP86, CYP96, and CYP72 family members were most abundant while possess only one to two members of CYP97, CYP716 and CYP722. Similarly, in *G. max* and *V. vinifera* where CYPs were distributed in 49 and 45 families respectively (Nelson et al. 2009), within A-type, majority members belongs to CYP71, CYP76, CYP81, and CYP82 families, while within non-A type, *G. max* possess higher number of members in CYP72, CYP90, CYP94 and *V. vinifera* possess higher CYP72, CYP86, and CYP716 family members. Moreover, no CYP710 members were present in *A. agallocha*,while four in *A. thaliana*, two in *G. max* and one in *V. vinifera* were reported (Jiu et al. 2020). Additionally, no members of CYP727 and CYP746 were found neither in *A. agallocha* nor in other three plants, except one in *G. max*. Probably during evolution in these plants, the members have lost or may evolved into other family. It was observed that distribution of number of CYPs in each family varies among the plant species which might indicate differences in their metabolites profile, adaptation to stress and developmental processes (Xu et al. 2015). Therefore, considering the association of agarwood with *Aquilaria* plant species, evolution of CYPs in them are possibly more selective towards defense and secondary metabolism. Additionally, only AaCYPs which belongs to CYP71, CYP85 and CYP86 clans (multiple-family clan) expanded through segmental (major event) and tandem duplication. This indicates that single-family clan members were less susceptible to duplication pressure while expansion of multiple-family clans in *A. agallocha* were majorly due to whole genome duplication events. Similarly, in *A. thaliana* and *G. max*, segmental or genome duplications were major driving force for CYPs expansion (Yu et al. 2017, Guttikonda et al., 2010), unlike in *V. vinifera*, where tandem duplications were major events (Jiu et al. 2020). The Ka/Ks < 0.05 of all the duplicated *AaCYPs* suggest their stabilizing selection during evolution (Yang et al. 2000). However, presence of orthologous pairs with all the 4 plant species, and few members with atleast one out of the four species suggest their involvement in speciation events and conserved functional role.

### 4.2. AaCYPs involvement in biosynthesis of important metabolites in *A. agallocha*

Pathway mapping and prediction of functional partners of AaCYPs revealed their possible functional role in developmental processes and biosynthesis of important metabolites reported in *Aquilaria* and agarwood (Li et al. 2021). The CYP51, CYP74, CYP86, and CYP97 families which remain conserved throughout Viridiplantae were present in *A. agallocha* suggesting their participation in the metabolite’s biosynthesis (Hansen et al. 2021). For instance, in lipids metabolism, CYP51G acts as 14α-demethylase in sitosterol biosynthesis (Lepesheva et al. 2006). In *A. thaliana*, CYP51G1 involves in demethylation of obtusifoliol (Kim et al. 2005). So, the only member AaCYP51G (82.1% sequence similarity with AtCYP51G1) may possibly involve in steroids biosynthesis including β-sitosterol identified in *Aquilaria*, and stigmasterol and ergosta-4,6,8(14),22-tetraen-3-one in agarwood (Li et al. 2021). Here, two members *viz*. AaCYP74A1 and AaCYP74A2 similar to *allene oxide synthase 1* (72.8% and 54.1%) of *S. lycopersicum* (Sivasankar et al. 2000) probably functions as AOSs in jasmonic acid biosynthesis. While, AaCYP74B1 which was similar to fatty acid hydroperoxide lyase (66.9%) probably involved in the biosynthesis of green leaf volatiles (GLVs) in *A. agallocha*. Interestingly, in *A. sinensis*, 9 GLVs including 2-hexanol, 2-ethyl-1-hexanol, (Z)-3-hexen-1-ol and so on were identified which were elevated upon biotic stress (Qiao et al. 2018) indicating CYP74B member’s defensive role in *Aquilaria*.

The CYP86 family members acts on fatty acids and contributes to the cell wall components like suberin, cutin and wax biosynthesis (Bak et al. 2011). In *A. agallocha*, AaCYP86A3 (78.8% similar with AtCYP86A1), AaCYP86A1 (76.8% with AtCYP86A8) and AaCYP86A2 (75.8% with PhCYP86A22) possibly participated in the cell wall biosynthesis. In *A. thaliana*, AtCYP86A1 and AtCYP86A8 catalyse omega-hydroxylation of saturated and unsaturated fatty acids (Benveniste et al. 1998, Wellesen et al. 2001) and, in *P. hybrida*, PhCYP86A22 also catalyses similar reaction in the biosynthesis of polyesters (Han et al. 2010). Interestingly, in *A. thaliana* and *G. barbadense*, CYP86 members also imparts resistance against plant pathogens (Xiao et al. 2004, Wang et al. 2020) which indicates multifunctional nature of plant CYP86s. Reports suggest threshold of approximately 50% similarity between the homologs below which the functional divergence increases (Sanger et al. 2007). Thus, the higher similarities of AaCYP86 members with these proteins illustrates their involvement not only in lipids biosynthesis but also possibly in defense in *A. agallocha* (Wilson et al. 2000).

The CYP97 family members are associated with carotenoid biosynthesis where they catalyse ring hydroxylation of carotenes in algae (Liang et al. 2020) and *A. thaliana* (AtCYP97A3 and AtCYP97C1) (Kim et al. 2009). The two members viz. AaCYP97A1 (77.5% similar to AtCYP97A3), and AaCYP97C1 (75.1% similar to AtCYP97C1) probably involves in carotenoid biosynthesis. In *A. thaliana*, CYP85, CYP90, and CYP724 family members involves in the regulation and biosynthesis of brassinosteroid (BRs), where AtCYP85A regulates BR C-6 oxidase expression (Castle et al. 2005); AtCYP90A1 and AtCYP90B1 in development (Choe et al. 2001, Bancos et al. 2002); AtCYP90C1 and AtCYP90D1 as C-23 hydroxylase (Ohnishi et al. 2006); while SlCYP90B3 and SlCYP724B2 acts as C-22 hydroxylase (Ohnishi et al. 2014). The *Aquilaria* members viz. AaCYP85A1 (similar clade with AtCYP85A1), AaCYP90C1 (71.1% match with AtCYP90C1), AaCYP90D1 (67.4% match with AtCYP90D1), and AaCYP724B1 (64.3% match with OsCYP724B1) probably participates in BR biosynthesis pathway and in plant immunity against pathogen (Wang et al. 2012, Tanabe et al. 2005).

Functionally versatile CYP71 clan is composite of diverse families as a result of gene duplication events (Bak et al. 2011) and lineage-specific proliferations (blooms) (Feyereisen 2011). The families CYP73, CYP84, and CYP98 involves in phenylpropanoid biosynthesis (Hansen et al. 2021). CYP73 members had been detected in *G. max* (CYP73A11) and *A. thaliana* (CYP73A5), and their functions were annotated based on similarity to CYP73A1 of *H. tuberosus* (Urban et al. 1994). In this study, two members namely AaCYP73A1 (86.5% similar to CYP73A11 of *G. max*) and AaCYP73A2 (82.1% similar to CYP73A100 of *P. ginseng*) possibly involved in conversion of cinnamate or cinnamoyl into coumaric acid or coumaroyl in *A. agallocha*. Additionally, CYP84 and CYP98 members involves in 5□ and 3□ hydroxylation of coniferaldehydes and coumaroyl-shikimate/coumaroyl-quinate, respectively (Humphreys et al. 1999, Mahesh et al. 2007). Interestingly *A. thaliana* member viz. AtCYP84A1 and AtCYP84A4 has 5 □ and 3 □-hydroxylase activity respectively towards p-coumaraldehyde (Weng et al. 2012), while AtCYP98A3 shows 3□-hydroxylase activity towards multiple substrates including p-coumarate, p-coumaroyl esters, and p-coumaraldehyde in the phenylpropanoid pathway (Schoch et al. 2001). The members, AaCYP84A1, AaCYP84A2 and AaCYP84A3 showed similarity in between 70.5 to 79.8% with AtCYP84A1, AaCYP98A1, and GmCYP98A2. Thus, AaCYP84 members probably utilizes substrates including ferulic acid, coniferylaldehyde and coniferylalcohol for 5□ hydroxylation, while AaCYP98A1 utilizes 4-coumaroylshikimate to generate caffeoyl or chlorogenic acids through 3□ hydroxylation. Although the above chemicals had been illustrated in *Aquilaria* (Li et al. 2021), no immediate or intermediate hydroxylated products have been mentioned in *Aquilaria* or agarwood till date. However, an end product named syringin which is derived from sinapyl alcohol has been reported (Li et al. 2021). Members of CYP75 family acts as flavonoids hydroxylase at 5□ and 3□ (CYP75B), and only 3□ (CYP75A) (Yonekura-Sakakibara et al. 2019). In *A. thaliana*, CYP75B1 catalyses the conversion of naringenin to eriodictyol and dihydrokaempferol to dihydroquercetin through 3□ hydroxylation (Schoenbohm et al. 2005), but no members of CYP75A subfamily were detected till date. In *A. agallocha*, AaCYP75A1 showed 60.9% similarity with CYP75A3 of *P. hybrida* whose function was assigned based on its paralog CYP75A1 (3□ and 5□ hydroxylase activity) (Shimada et al. 1999). The three members AaCYP75B1, AaCYP75B2, and AaCYP75B3 with 67.9%, 57.9%, and 64.3% similarity to AtCYP75B1 indicates similar substrate catalysis. Hence, identification of flavonoids viz. kaempferol, luteolin, sakuranetin, and quercetin in *Aquilaria* (Li et al. 2021) indicates collective role of CYP73, CYP75 and CYP98 family members. However, few AaCYPs had low sequence similarity (29 to 43%) with characterized CYPs, possibly such enzymes may have evolved in *Aquilaria* for novel substrate and product profile with rare biochemical properties. Nevertheless, the AaCYP members found associated with defense compounds including jasmonic acids and brassinosteroid, and pharmaceutically important metabolites such as phenolics and flavonoids illustrates their functional role and complex regulation in *A. agallocha* during biotic and abiotic stress.

### 4.3. Potential role of AaCYPs in biosynthesis of fragrant agarwood constituents in *A. agallocha*

Sesquiterpenoids and 2-(2-phenylethyl) chromones (PECs) are the vital constituents in agarwood which imparts its unique odor (Chen et al. 2012, Naef 2011, Yagura et al. 2003). Till date, more than 180 sesquiterpenoids, 200 2-(2-phenyl) chromones and 10 sesquiterpenoid-chromone derivatives has been identified in agarwood (Li et al. 2021, Chen et al. 2012). As per previous reports, the members CYP71AV, CYP71BA, CYP71BL, CYP71D, and CYP71BE involved in sesquiterpenoid biosynthesis (Hamberger et al. 2013). In *A. agallocha*, the two members viz. AaCYP71D7, and AaCYP71D17 with 52.8% and 48.7% sequence similarity to CYP71D55 of *H. muticus* possibly associates with sesquiterpenoids biosynthesis. CYP71D55 shows activity towards 5 sesquiterpenes namely, valencene, vestipiradiene, 4-epi-eremophilene, 5-epiaristolochene, and cedr-8-ene which generates sesquiterpenoids nootkatone, solavetivol/solavetivone, 4-epieremophilen-2beta-ol, 2beta-hydroxy-5-epiaristolochene, and cedr-8-en-15-ol, respectively (Takahashi et al. 2007). The expression of AaCYP71D7 was insignificant in the RNA-seq data, but elevation of AaCYP71D17 (3.05 log2FC) indicates its probable involvement in generation of sesquiterpenoids and their derivatives such as (+)-trans-nootkatol, 11-hydroxy-valenc-1(l0)-en-2-on; valenca-1(10),8-dien-11-ol; vetispira-2(11),6(14)-dien-7-ol; and eremophil-9-ene-11,12,13-triol etc. which were previously detected in agarwood (Li et al. 2021). Moreover, promiscuous nature of CYPs has further facilitate diversification in terms of metabolites profile (Hamberger et al. 2013). For instance, germacrene A oxidases (GAOs), not only catalyses germacrene A into germacrene A acid, but also accepts other sesquiterpenes such as amorphadiene and germacrene D (Eljounaidi et al. 2014). Similarly, CYP71D subfamily members also accepts monoterpenes (Wust et al. 2001) and diterpenes (Wang et al. 2003) besides sesquiterpenes (Ralston et al. 2001). In this study, the members AaCYP71D1, AaCYP71D4, AaCYP71D10, and AaCYP71D11 whose expression were elevated up to 5-10 log2FC in RNA-seq data, showed similarity in the range of 49-53% with CYP71D10 (involves in monoterpenoid biosynthesis) of *G. max* (Siminszky et al. 1999). Additionally, two members AaCYP71D18 and AaCYP71D19 elevated up to 8-9 logFC similar to salviol synthase (involves in diterpenoid biosynthesis) of *S. pomifera* (Ignea et al. 2016) in the range of 45-48%. Considering the promiscuity of CYP450s, these AaCYP members may also show activity towards sesquiterpenes besides mono and diterpenes. However, in this regard biochemical characterization would be an interesting opportunity for further research. The sweet and lasting scent of agarwood is believed to be because of 2-(2-phenylethyl)chromones (PECs), but mechanism for their biosynthesis is still not clear, except for characterization of few polyketide synthases in *A. sinensis* which biosynthesizes the precursors of PECs (Wang et al. 2022, Wang et al. 2017). Interestingly, the core structures of the PECs detected in *Aquilaria* (Liao et al. 2018) suggest that it derives from the common phenylpropanoid pathway (Borejsza-Wysocki et al. 1994) through polyketide synthases (type III PKS) (Wang et al. 2016) and subsequent hydroxylation and *O*-methylation producing highly oxygenated chromones (Wang et al. 2022, Liao et al. 2018). Therefore, we assume that the oxidation and subsequent modifications into different types of chromones (Li et al. 2021) are probably catalyses through monooxygenase activity of *Aquilaria* CYPs. This study could only detect two CYP73A subfamily members (AaCYP73A1 and AaCYP73A2) that possibly act as cinnamate monooxygenase and generates the precursors (p-coumaroyl CoA and cinnamoyl CoA) of phenolics, flavonoids and agarwood PECs. Taking into account the elevation of *AaCYP73A2* in RNA-seq and qRT-PCR expression data, and no information regarding the same in PEC biosynthesis in *Aquilaria* or in any other plants till date, it will be worth experimenting to assess if these members could take part in the biosynthesis of PECs in *A. agallocha*.

Additionally, validation of the transcript abundance of the candidate members in phenylpropanoid and sesquiterpenoid biosynthesis in infected wood tissue of *A. agallocha* illustrates their participation in plant defense, biosynthesis of crucial *Aquilaria* chemicals and also probably during agarwood formation. It is worth mentioning that the members viz. *AaCYP71D7, AaCYP71D10*, and *AaCYP71D17* with higher expression only in infected wood tissue but not in methyl jasmonate treated callus, indicates their tissue specific nature or involvement in decoration of sesquiterpenes specifically present in agarwood. The other three genes (*AaCYP71D1*, *AaCYP71D4*,and *AaCYP71D11*) which were elevated both in MeJ treated callus and infected wood of *A. agallocha* may also contribute in sesquiterpenoid biosynthesis both in agarwood and stress induced callus. This correlates with previous reports that MeJ can induce production of sesquiterpenoids (Kumeta and Ito 2010) and 2-(2-phenylethyl) chromones in callus of *A. sinensis* (Dong et al. 2018). Hence, the CYPs could be exploited as candidate members to understand agarwood formation *in-vitro* as well.

## 5. Conclusion

This study provides the comprehensive overview of *A. agallocha’*s CYP450 superfamily, including protein identification, characterization, phylogenetic position, conserved motifs, evolution, functional classification, as well as in-silico and experimental expression during stress, which collectively collate important features of the family members in *Aquilaria* plant species. Additionally, provides insight into the possible candidate CYP members and their functional position in the biosynthesis of valuable metabolites identified earlier in agarwood and *Aquilaria* plant species. Although the actual molecular mechanism of quality agarwood formation is still a mystery, the metabolites profile of agarwood firmly asserts CYP450s involvement, and evidences for the same has been outlined comprehensively in the study. In future, biochemical characterization of the candidate members will not only facilitate in-vitro production of crucial natural metabolites of *Aquilaria* plants but also help better understand agarwood formation.

## Data availability

The data supporting the finding of this study is provided in the manuscript and its supplementary material.

## Ethics approval

Not applicable

## Acknowledgments

The authors are indebted to Gauhati University for providing the technical facility. The authors also acknowledge Department of Biotechnology (DBT), Government of India for providing the financial aid.

## Abbreviations

CYP: Cytochrome P450
AaCYP: *Aquilaria agallocha* Cytochrome P450
PEC: Phenylethyl chromones
DMSO: Dimethyl sulfoxide
HMM: Hidden Markov model
CTAB: Cetyltrimethyl ammonium bromide
GO: Gene ontology
KEGG: Kyoto Encyclopaedia of Genes and Genomes database

## Supplementary files

**Additional file 1.** General properties of AaCYPs and their deduced protein sequences identified in the genome of *Aquilaria agallocha*.

**Additional file 2.** Distribution of AaCYPs in 8 CYP clans.

**Additional file 3.** Cis Regulatory elements in the 2000bp promoter region of 136 *AaCYPs*.

**Additional file 4.** Orthologous gene pairs of *Aquilaria agallocha CYPs* with *Vitis vinifera, Arabidopsis thaliana, Glycine max* and *Solanum lycopersicum CYPs*.

**Additional file 5.** Duplicated *Aquilaria agallocha CYP* gene pairs.

**Additional file 6.** Ka, Ks, and MYA of duplicated gene pairs.

**Additional file 7.** Functional grouping of AaCYPs based on GO terms.

**Additional file 8.** Mapping of AaCYPs in KEGG pathway database for functional prediction and categorization.

**Additional file 9.** Gene expression of significantly regulated genes and the pathways involvement based on KEGG, Uniprot and literature.

**Additional file 10.** Details of primers and PCR carried out in the present study.

**Additional file 11.** Evaluation of 3D structure of 8 members of respective clans. A) ProQ2 quality assessment. B) Clashes. C) Rotamers. D) Ramachandran analysis. E) Alignment confidence. F) conservation. G) Pocket detection

**Additional file 12.** The four conserve motifs i.e., heme-binding (violet colour loop), PxRx (blue colour helix), K-helix (cyan colour helix), and I-helix (pink colour helix) indicated in the models.

## References

1. A. Butrón, Y.C. Chen, G.E. Rottinghaus, et al., Genetic variation at bx1 controls DIMBOA content in maize. Theor. Appl. Genet. 120 2010 721–734.

2. A. Das, K. Begum, S. Akhtar, et al., Genome-wide detection and classification of terpene synthase genes in Aquilaria agallochum, Physiol. Mol. Biol. Plants 27 2021 1711–1729.

3. A. Krokida, C. Delis, K. Geisler, et al., A metabolic gene cluster in Lotus japonicus discloses novel enzyme functions and products in triterpene biosynthesis. New Phytologist 200 2013 675–690.

4. A. Rai, R. Singh, P.A. Shirke, et al., Expression of rice CYP450-like gene (os08g01480) in arabidopsis modulates regulatory network leading to heavy metal and other abiotic stress tolerance, PLoS ONE 2015, 10, e0138574.

5. A. Ray, E. Lindahl, B. Wallner, Improved model quality assessment using ProQ2, BMC Bioinformatics 13 2012 224.

6. B. Hamberger, S. Bak, Plant P450s as versatile drivers for evolution of species-specific chemical diversity, Philos. Trans. R. Soc. Lond. B Biol. Sci. 368(1612) 2013.

7. B. Hu, J. Jin, A.Y. Guo, et al., GSDS 2.0: an upgraded gene feature visualization server, Bioinformatics 31(8) 2015 1296–1297.

8. B. Ma, Y. Luo, L. Jia, et al., Genome wide identification and expression analyses of cytochrome P450 genes in mulberry (Morus notabilis), J. Integr. Plant Biol. 56 2014 887–901.

9. B. Siminszky, F.T. Corbin, E.R. Ward, et al., Expression of a soybean cytochrome P450 monooxygenase cDNA in yeast and tobacco enhances the metabolism of phenylurea herbicides, Proc. Natl. Acad. Sci. U.S.A. 96(4) 1999 1750–1755.

10. C. Chen, H. Chen, Y. Zhang, et al., TBtools: An Integrative Toolkit Developed for Interactive Analyses of Big Biological Data. Mol Plant. 13(8) 2020 1194–1202.

11. C. Ignea, A. Athanasakoglou, E. Ioannou, et al., Carnosic acid biosynthesis elucidated by a synthetic biology platform. Proc. Natl. Acad. Sci. U.S.A. 113(13) 2016 3681–3686.

12. C. Lopez-Ortiz, S.K. Dutta, P. Natarajan, et al., Genome-wide identification and gene expression pattern of ABC transporter gene family in Capsicum spp, PLoS One 14 2019 e0215901.

13. C. Wasternack, M. Strnad, Jasmonate signaling in plant stress responses and development-Active and inactive compounds, New Biotechnol. 33 2016 604–613.

14. C.A. Wilson, J. Kreychman, M. Gerstein, Assessing annotation transfer for genomics: quantifying the relations between protein sequence, structure and function through traditional and probabilistic scores, J. Mol. Biol. 297 2000 233–249.

15. C.C. Hansen, D.R. Nelson, B.L. Møller, et al., Plant cytochrome P450 plasticity and evolution, Molecular Plant 14(8) 2021 1244–1265.

16. C.J.A. Sigrist, L. Cerutti, N. Hulo, et al., PROSITE: a documented database using patterns and profiles as motif descriptors, Brief Bioinform. 3(3) 2002 265–74.

17. C.S. Tan, N.M. Isa, I. Ismail, et al., Agarwood Induction: Current Developments and Future Perspectives. Front. Plant Sci. 10 2019 122.

18. D. Kim, B. Langmead, S. Salzberg, HISAT: a fast-spliced aligner with low memory requirements, Nat. Methods 12 2015 357–360.

19. D. Nelson, D. Werck-Reichhart, A P450-centric view of plant evolution, Plant J. 66(1) 2011 194–211.

20. D. Qin, H. Wu, H. Peng, et al., Heat stress-responsive transcriptome analysis in heat susceptible and tolerant wheat (Triticum aestivum L.) by using Wheat genome array, BMC Genom. 9 2008 432.

21. D. Wang, Y. Zhang, Z. Zhang, et al., KaKs_calculator 2.0: a toolkit incorporating gamma-series methods and sliding window strategies, Genomics Proteomics Bioinformatics 8 2010 77–80.

22. D. Werck-Reichhart, R. Feyereisen, Cytochromes P450: a success story, Genome Biol. 1 2000 1–9.

23. D.R. Nelson, Cytochrome P450 and the individuality of species, Arch. Biochem. Biophys. 369(1) 1999 1–100.

24. D.R. Nelson, Cytochrome P450 diversity in the tree of life, Biochim Biophys Acta Proteins Proteom. 1866(1) 2018 141–154.

25. D.R. Nelson, Cytochrome P450 genes from the sacred Lotus genome, Trop Plant Biol. 6 2013 138–51.

26. D.R. Nelson, L. Koymans, T. Kamataki, et al., P450 superfamily: Update on new sequences, gene mapping, accession numbers and nomenclature, Pharmacogenetics 6 1996 1–42.

27. D.R. Nelson, R. Ming, M. Alam, et al., Comparison of Cytochrome P450 Genes from Six Plant Genomes, Tropical Plant Biol. 2008 1 216–235.

28. D.R. Nelson, The Cytochrome P450 Homepage, Human Genomics 4 2009 59.

29. D.T. Ahmaed, A.D. Kulkarni, Sesquiterpenes and Chromones of Agarwood: A Review, Malays. J. Chem. 19(1) 2017 33–58.

30. E. Wang, G.J. Wagner, Elucidation of the functions of genes central to diterpene metabolism in tobacco trichomes using posttranscriptional gene silencing, Planta 216 2003 686–691.

31. F. Pinot, F. Beisson, Cytochrome P450 metabolizing fatty acids in plants: characterization and physiological roles. FEBS J. 278(2) 2011 195–205.

32. F. Xiao, S.M. Goodwin, Y. Xiao, et al., Arabidopsis CYP86A2 represses Pseudomonas syringae type III genes and is required for cuticle development, EMBO J. 23(14) 2004 2903–2913.

33. F.P. Guengerich, A history of the roles of cytochrome P450 enzymes in the toxicity of drugs, Toxicol. Res. 37 2021 1–23.

34. G Liao, W.H Dong, J.L. Yang, et al., Monitoring the Chemical Profile in Agarwood Formation within One Year and Speculating on the Biosynthesis of 2-(2-Phenylethyl) Chromones, Molecules 23(6) 2018 1261.

35. G.I. Lepesheva, M.R. Waterman, Sterol 14alpha-demethylase cytochrome P450 (CYP51), a P450 in all biological kingdoms. Biochim. Biophys. Acta. 1770(3) 2007 467–477.

36. H. Qiao, P. Lu, S. Liu, et al., Volatiles from Aquilaria sinensis damaged by Heortia vitessoides larvae deter the conspecific gravid adults and attract its predator Cantheconidea concinna, Scientific reports, 8(1) 2018 15067

37. H. Wang, Q. Wang, Y. Liu, et al., PCPD: Plant cytochrome P450 database and web-based tools for structural construction and ligand docking, Synth. Syst. Biotechnol. 6(2) 2021 102–109.

38. H.B. Kim, H. Schaller, C.H. Goh, et al., Arabidopsis cyp51 mutant shows postembryonic seedling lethality associated with lack of membrane integrity. Plant Physiol. 138(4) 2005 2033–2047.

39. H.Q. Chen, J.H. Wei, J.S. Yang, et al., Chemical constituents of agarwood originating from the endemic genus Aquilaria plants, Chem Biodivers. 9 2012 236–250.

40. I. Benveniste, N. Tijet, F. Adas, et al., CYP86A1 from Arabidopsis thaliana encodes a cytochrome P450-dependent fatty acid omega-hydroxylase, Biochem Biophys Res Commun. 243(3) 1998 688–693.

41. I.G. Denisov, T.M. Makris, S.G. Sligar, et al., Structure and chemistry of cytochrome P450, Chem. Rev. 105(6) 2005 2253–2277.

42. J. Booker, T. Sieberer, W. Wright, et al., MAX1 encodes a cytochrome P450 family member that acts downstream of MAX3/4 to produce a carotenoid-derived branch-inhibiting hormone, Developmental Cell 8 2005 443–449.

43. J. Castle, M. Szekeres, G. Jenkins, et al., Unique and overlapping expression patterns of Arabidopsis CYP85 genes involved in brassinosteroid C-6 oxidation, Plant Molecular Biology 57 2005 129–140.

44. J. Chappell, R.M. Coates, Sesquiterpenes, Comprehensive Natural Products II, 1 2010 609–641 ISBN 9780080453828.

45. J. Han, J.M. Clement, J. Li, et al., The Cytochrome P450 CYP86A22 Is a Fatty Acyl-CoA ω-Hydroxylase Essential for Estolide Synthesis in the Stigma of Petunia hybrida2. Journal of Biological Chemistry 285(6) 2010 2986–3996.

46. J. Kim, J.J. Smith, L. Tian, et al., The Evolution and Function of Carotenoid Hydroxylases in Arabidopsis, Plant and Cell Physiology 50(3) 2009 463–479.

47. J. Xu, X.Y Wang, W.Z Guo, The cytochrome P450 superfamily: Key players in plant development and defense, Journal of Integrative Agriculture 14(9) 2015 1673–1686.

48. J. Yu, S. Tehrim, L. Wang, et al., Evolutionary history and functional divergence of the cytochrome P450 gene superfamily between Arabidopsis thaliana and Brassica species uncover effects of whole genome and tandem duplications, BMC Genomics 18 2017 733.

49. J.E. Kim, K.M. Cheng, N.E. Craft, et al., Overexpression of Arabidopsis thaliana carotenoid hydroxylases individually and in combination with a β-carotene ketolase provides insight into in vivo functions, Phytochemistry 71 2010 168–178.

50. J.K. Weng, Y. Li, H. Mo, et al., Assembly of an Evolutionarily New Pathway for α-Pyrone Biosynthesis in Arabidopsis, SCIENCE 337(6097) 2012 960–964.

51. J.M. Humphreys, M.R. Hemm, C. Chapple, New routes for lignin biosynthesis defined by biochemical characterization of recombinant ferulate 5-hydroxylase, a multifunctional cytochrome P450-dependent monooxygenase, Proc. Natl. Acad. Sci. U.S.A. 96(18) 1999 10045–10050.

52. K. Wei, H. Chen, Global identification, structural analysis and expression characterization of cytochrome P450 monooxygenase superfamily in rice, BMC Genomics 19 2018 35.

53. K. Wellesen, F. Durst, F. Pinot, et al., Functional analysis of the LACERATA gene of Arabidopsis provides evidence for different roles of fatty acid omega -hydroxylation in development, Proc. Natl. Acad. Sci. U.S.A. 98(17) 2001 9694–9699.

54. K. Yonekura-Sakakibara, Y. Higashi, R. Nakabayashi, The Origin and Evolution of Plant Flavonoid Metabolism, Front. Plant Sci. 10 2019 943.

55. L. Li, H. Cheng, J. Gai, et al., Genome-wide identification and characterization of putative cytochrome P450 genes in the model legume Medicago truncatula, Planta. 226(1) 2007 109–123.

56. L. Peng, Y. Zhao, H. Wang, et al., Functional Study of Cytochrome P450 Enzymes from the Brown Planthopper (Nilaparvata lugens Stål) to Analyze Its Adaptation to BPH-Resistant Rice, Front. Physiol. 8 2017 972.

57. L. Ralston, S.T. Kwon, M. Schoenbeck, et al., Cloning, heterologous expression, and functional characterization of 5-epi-aristolochene-1,3-dihydroxylase from tobacco (Nicotiana tabacum), Arch Biochem Biophys. 393(2) 2001 222–235.

58. L.A. Kelley, M.J.E. Sternberg, Protein structure prediction on the Web: a case study using the Phyre server, Nat. Protoc. 4 2009 363–371.

59. M. Gao, X. Han, Y. Sun, et al., Overview of sesquiterpenes and chromones of agarwood originating from four main species of the genus Aquilaria, RSC Adv. 9 2019 4113–4130.

60. M. Lescot, P. Déhais, G. Thijs, et al., PlantCARE, a database of plant cis-acting regulatory elements and a portal to tools for in silico analysis of promoter sequences, Nucleic Acids Res. 30 2002 325–327.

61. M. Liu, Z. Ma, A. Wang, et al., Genome-wide investigation of the auxin response factor gene family in Tartary buckwheat (Fagopyrum tataricum), Int. J. Mol. Sci. 19 2018 3526.

62. M. Mizutani, D. Ohta, Diversification of P450 genes during land plant evolution, Annu. Rev. Plant Biol. 61 2010 291–315.

63. M. Mizutani, D. Ohta, Diversification of P450 genes during land plant evolution. Annu. Rev. Plant Biol. 61 2010 291–315.

64. M. Tamiru, J.R. Undan, H. Takagi, et al., A cytochrome P450, OsDSS1, is involved in growth and drought stress responses in Rice (Oryza sativa L.), Plant Mol Biol. 88(1-2) 2015 85–99.

65. M. Wüst, D.B. Little, M. Schalk, et al., Hydroxylation of Limonene Enantiomers and Analogs by Recombinant (-)-Limonene 3-and 6-Hydroxylases from Mint (Mentha) Species: Evidence for Catalysis within Sterically Constrained Active Sites, Archives of Biochemistry and Biophysics Volume 387(1) 2001 125–136.

66. M.A. Schuler, The role of cytochrome P450 monooxygenases in plant-insect interactions, Plant Physiol. 112 1996 1411–1419.

67. M.H. Liang, H. Xie, H.H. Chen, et al., Functional Identification of Two Types of Carotene Hydroxylases from the Green Alga Dunaliella bardawil Rich in Lutein. ACS Synth Biol. 9(6) 2020 1246–1253.

68. M.I. Love, W. Huber, S. Anders, Moderated estimation of fold change and dispersion for RNA-seq data with DESeq2, Genome Biology 15(12) 2014 550.

69. M.M. Xie, D.P. Gong, F.X. Li, et al., Genome-wide analysis of cytochrome P450 monooxygenase genes in the tobacco, Hereditas 35(3) 2013 379–387.

70. M.N. Wass, L.A. Kelley, M.J. Sternberg, 3DLigandSite: predicting ligand-binding sites using similar structures, Nucleic Acids Res. 38 2010 W469–W473.

71. M.R. Islam, B.S Bhau, S. Banu, Gene expression analysis associated with agarwood formation in Aquilaria malaccensis. Plant Physiol. Rep. 25 2020 304–314.

72. M.R. Islam, S. Banu, An improved cost-effective method of RNA extraction from Aquilaria malaccensis, Acta Scientific Agriculture 3(2) 2019 30–38.

73. M.R. Islam, S. Banu, Transcript profiling leads to biomarker identification for agarwood resin-loaded Aquilaria malaccensis, Trees 35 2021 2119–2132.

74. M.R. Islam, C. Chakraborty, S. Banu, Isolation and Characterization of Bacteria and Fungi Associated with Agarwood Fermentation, Current Microbiology 79 2022.

75. M.S. Kumar, P.R. Babu, K.V. Rao, et al., Organization and classification of cytochrome p450 genes in castor (Ricinus communis L.), Proc. Natl. Acad. Sci. India Sect. B Biol. Sci. 84 2014 131–143.

76. P. Wang, L. Ma, Y. Li, et al., Transcriptome analysis reveals sunflower cytochrome P450 CYP93A1 responses to high salinity treatment at the seedling stage, Genes Genom. 39 2017 581–591.

77. P.R. Babu, K.V. Rao, V.D. Reddy, Structural organization and classification of cytochrome P450 genes in flax (Linum usitatissimum L.), Gene 513(1) 2013 156–162.

78. R. Bari, J.D.G. Jones, Role of plant hormones in plant defence responses. Plant Mol. Biol. 69 2009 473–488.

79. R. Feyereisen, Arthropod CYPomes illustrate the tempo and mode in P450 evolution, Biochimica et Biophysica Acta (BBA) - Proteins and Proteomics, 1814(1) 2011 19–28.

80. R. Mohamed, P.L. Jong, A.K. Kamziah, Fungal inoculation induces agarwood in young Aquilaria malaccensis trees in the nursery. J. For. Res. 25 2014 201–204.

81. R. Naef, The volatile and semi-volatile constituents of agarwood, the infected heartwood of Aquilaria species: a review, Flavour Fragr J. 26 2011 73–87.

82. R.C. Misra, S. Sharma, Sandeep, et al., Two CYP716A subfamily cytochrome P450 monooxygenases of sweet basil play similar but nonredundant roles in ursane-and oleananetype pentacyclic triterpene biosynthesis, New Phytologist 214 2017 706–720.

83. R.K. Hughes, S.D. Domenico, A. Santino, Plant cytochrome CYP74 family: biochemical features, endocellular localisation, activation mechanism in plant defence and improvements for industrial applications, ChemBioChem 10 2009 1122–1133.

84. R.S. Li, J.H. Zhu, D. Guo, et al., Genome-wide identification and expression analysis of terpene synthase gene family in *Aquilaria sinensis*, Plant Physiology and Biochemistry 164 2021 185–194.

85. S. Anders, W. Huber, Differential expression analysis for sequence count data. Genome Biol. 11 2010, R106.

86. S. Bak, F. Beisson, G. Bishop, et al., Cytochromes p450, Arabidopsis Book 2011(9) 2011.

87. S. Bancos, T. Nomura, T. Sato, et al., Regulation of transcript levels of the Arabidopsis cytochrome p450 genes involved in brassinosteroid biosynthesis, Plant Physiology 130 2002 504–513.

88. S. Choe, S. Fujioka, T. Noguchi, et al., Overexpression of DWARF4 in the brassinosteroid biosynthetic pathway results in increased vegetative growth and seed yield in Arabidopsis, The Plant Journal 2001 26 573–582.

89. S. Graham-Lorence, B. Amarneh, R.E. White, et al., A three-dimensional model of aromatase cytochrome P450, Protein Sci. 4(6) 1995 1065–1080.

90. S. Guillaume, G. Simon, M. Morant, et al., CYP98A3 from Arabidopsis thaliana Is a 3’-Hydroxylase of Phenolic Esters, a Missing Link in the Phenylpropanoid Pathway, Journal of Biological Chemistry 276(39) 2001 36566–36574.

91. S. Islam, S.D. Sajib, Z.S. Jui, et al., Genome-wide identification of glutathione S-transferase gene family in pepper, its classification, and expression profiling under different anatomical and environmental conditions, Sci. Rep. 9 2019 1–15.

92. S. Jiu, Y. Xu, J. Wang, et al., The Cytochrome P450 Monooxygenase Inventory of Grapevine (Vitis vinifera L.): Genome-Wide Identification, Evolutionary Characterization and Expression Analysis, Frontiers in Genetics 11 2020 44.

93. S. Kovaka, A.V. Zimin, G.M. Pertea, et al., Transcriptome assembly from long-read RNA-seq alignments with StringTie2, Genome Biol. 20(1) 2019.

94. S. Kumar, G. Stecher, M. Li, et al., MEGA X: molecular evolutionary genetics analysis across computing platforms, Mol. Biol. Evol. 35(6) 2018 1547–1549.

95. S. Schoenbohm, S. Martens, C. Eder, et al., Identification of the Arabidopsis thaliana Flavonoid 3’-Hydroxylase Gene and Functional Expression of the Encoded P450 Enzyme, Biol. Chem. 381(8) 2000 749–753.

96. S. Sivasankar, B. Sheldrick, S.J. Rothstein, Expression of Allene Oxide Synthase Determines Defense Gene Activation in Tomato, Plant Physiology 122(4) 2000 1335–1342.

97. S. Takahashi, Y.S. Yeo, Y. Zhao, et al., Functional Characterization of Premnaspirodiene Oxygenase, a Cytochrome P450 Catalyzing Regio-and Stereo-specific Hydroxylations of Diverse Sesquiterpene Substrates, Journal of Biological Chemistry 282(43) 2007 31744–31754.

98. S. Wang, Z. Yu, C. Wang, et al., Chemical Constituents and Pharmacological Activity of Agarwood and Aquilaria Plants, Molecules 23(2) 2018 342.

99. S.C. Potter, A. Luciani, S.R. Eddy, et al., HMMER web server: 2018 update. Nucleic Acids Res. 46(W1) 2018 W200–4.

100. S.K. Guttikonda, J. Trupti, N.C. Bisht, et al., Whole genome co-expression analysis of soybean cytochrome P450 genes identifies nodulation-specific P450 monooxygenases, BMC Plant Biol. 10(1) 2010 243.

101. S. Möller, M.D.R. Croning, R. Apweiler, Evaluation of methods for the prediction of membrane spanning regions, Bioinformatics 17(7) 2001 646–653.

102. S.M. Paquette, S. Bak, R. Feyereisen, Intron–exon organization and phylogeny in a large superfamily, the paralogous cytochrome P450 genes of Arabidopsis thaliana, DNA Cell Biol. 19 2000 307–317.

103. S.X. Ge, D. Jung, R. Yao. ShinyGO: a graphical gene-set enrichment tool for animals and plants, Bioinformatics 36(8) 2020 2628–2629.

104. T. Morikawa, H. Saga, H. Hashizume, et al., CYP710A genes encoding sterol C22-desaturase in Physcomitrella patens as molecular evidence for the evolutionary conservation of a sterol biosynthetic pathway in plants, Planta 229 2009 1311–1322.

105. T. Moses, J. Pollier, Q. Shen, et al., OSC2 and CYP716A14v2 catalyze the biosynthesis of triterpenoids for the cuticle of aerial organs of Artemisia annua, The Plant Cell 27 2015 286–301.

106. T. Ohnishi, B. Watanabe, K. Sakata, et al., CYP724B2 and CYP90B3 function in the early C-22 hydroxylation steps of brassinosteroid biosynthetic pathway in tomato, Bioscience Biotechnology and Biochemistry, 70 2014 2071–2080.

107. T. Ohnishi, A.M. Szatmari, B. Watanabe, et al., C-23 Hydroxylation by Arabidopsis CYP90C1 and CYP90D1 Reveals a Novel Shortcut in Brassinosteroid Biosynthesis, The Plant Cell 18(11) 2006 3275–3288.

108. T. Yagura, M. Ito, F. Kiuchi, et al., Four new 2-(2-phenylethyl) chromone derivatives from withered wood of Aquilaria sinensis, Chem. Pharm. Bull (tokyo) 51(5) 2003 560–564.

109. U. Subasinghe, D. Hettiarachchi, Agarwood resin production and resin quality of Gyrinops walla, Gaertn. Int. J. Agr. Sci. 3 2013 357–362.

110. V. Mahesh, R. Million-Rousseau, P. Ullmann. N. Chabrillange, et al., Functional characterization of two p-coumaroyl ester 3’-hydroxylase genes from coffee tree: evidence of a candidate for chlorogenic acid biosynthesis, Plant Mol Biol. 64 2007 145–159.

111. V. Sangar, D.J. Blankenberg, N. Altman, et al., Quantitative sequence-function relationships in proteins based on gene ontology, BMC Bioinformatics 8 2007 294.

112. W. Li, Q.H. Chen, H. Wang, et al., Natural products in agarwood and Aquilaria plants: chemistry, biological activities and biosynthesis, Nat Prod Rep. 38 2021 528–556.

113. W. Sun, Z. Ma, L. Liu, Cytochrome P450 family: Genome-wide identification provides insights into the rutin synthesis pathway in Tartary buckwheat and the improvement of agricultural product quality, International Journal of Biological Macromolecules 164 2020 4032–4045.

114. W. Xu, S. Bak, A. Decker, et al., Microarray-based analysis of gene expression in very large gene families: the cytochrome P450 gene superfamily of Arabidopsis thaliana, Gene 272(1-2) 2001 61–74.

115. X. Dong, B. Gao, Y. Feng, et al., Production of 2-(2-phenylethyl) chromones in Aquilaria sinensis calli under different treatments, Plant Cell Tiss. Organ Cult. 135 2018 53–62.

116. X. Li, J.B. Zhang, B. Song, et al., Resistance to Fusarium head blight and seedling blight in wheat is associated with activation of a cytochrome P450 gene, Phytopathology 2010 100 183–191.

117. X. Liu, X. Zhu, H. Wang, et al., Discovery and modification of cytochrome P450 for plant natural products biosynthesis, Synth. Syst. Biotechnol. 5(3) 2020 187–199.

118. X.H. Wang, B.W. Gao, Y. Nakashima, et al., Identification of a diarylpentanoid-producing polyketide synthase revealing an unusual biosynthetic pathway of 2-(2-phenylethyl) chromones in agarwood, Nat Commun. 13 2022 348.

119. X. Wang, Z. Zhang, X. Dong, et al., Identification and functional characterization of three type III polyketide synthases from Aquilaria sinensis calli, Biochemical and Biophysical Research Communications. 486(4) 2017 1040–1047.

120. Y. Liu, H. Chen, Y. Yang, et al., Whole tree agarwood-inducing technique: an efficient novel technique for producing high-quality agarwood in cultivated Aquilaria sinensis trees, Molecules 18 2013 3086–3106.

121. Y. Okudera, M. Ito, Production of agarwood fragrant constituents in Aquilaria calli and cell suspension cultures. Plant Biotechnology 26(3) 2009 307–315.

122. Y. Shimada, R. Nakano-Shimada, M. Y. Okinaki, et al., Expression of chimeric P450 genes encoding flavonoid-3’,5’-hydroxylase in transgenic tobacco and petunia plants, FEBS Letters 461(3) 1999 241–245.

123. Y. Wang, H. Tang, J.D. DeBarry, et al., {MCScanX}: a toolkit for detection and evolutionary analysis of gene synteny and collinearity, Nucleic Acids Res. 40(7) 2012 e49.

124. Y.Y. Liu, J.H. Wei, Z.H. Gao, et al., A review of quality assessment and grading for agarwood, Chin. Herb. Med. 9 2017 22–30.

125. Z. Yang, J.P. Bielawski, Statistical methods for detecting molecular adaptation, Trends in Ecology & Evolution 15(12) 2000 496–503.

126. Z.Y. Wang, Brassinosteroids modulate plant immunity at multiple levels, Proc. Natl. Acad. Sci. U.S.A. 109(1) 2012 7–8.

