## Supplementary material for "Genome-wide investigation of Cytochrome P450 superfamily of *Aquilaria agallocha*: association with terpenoids and phenylpropanoids biosynthesis": Additional file 10.pdf

### Additional file 9: Details of primers and PCR carried out in the present study

1. *Table: Primer sequences used for gene expression analysis (RT-PCR) in the study*

| Genes | Primer sequence (5′-3′) | Ta(˚C) | Predicted pathway |
|---|---|---|---|
| *AaCYP73A2* | ATCGAGGCCACCAAGTTTAAT | 54 | Phenylpropanoid |
| | GCACCATTAGCAGCCATTATTT | | |
| *AaCYP84A1* | CCATCATAGACGACCACATGAA | 57 | Phenylpropanoid |
| | CCCTCCAAACATCACATCCA | | |
| *AaCYP98A1* | GGATAGAGCATGTTGTGGGATAG | 57 | Phenylpropanoid |
| | GGGTCATTTGTTGGATGTTGATG | | |
| *AaCYP84A2* | ATCATTGACGAGCACATGGA | 57 | Phenylpropanoid |
| | GTTTCTGTCCCTCCGAACAT | | |
| *AaCYP71D11* | GATCTCCAAATGCCCTACTTG | 56 | Terpenoid |
| | ACGTCTTGGCCAGATCCCTC | | |
| *AaCYP71D4* | ACGACCTTCTTGAATGCTTCCC | 60 | Terpenoid |
| | GTCGGTGAGGCATATCACCCAT | | |
| *AaCYP71D1* | GAGCAGCAATTCTTTTCATCTC | 59 | Terpenoid |
| | AGGTCATGATGTGGCAGACCTTT | | |
| *AaCYP71D7* | CTTTCCCAGATAATCGTTTC | 50 | Terpenoid |
| | AGACGGATAAAGATCTGC | | |
| *AaCYP71D17* | AACCCTAAGGTTGCACCCTA | 55 | Terpenoid |
| | GATCCCTAACGATTGCCCAC | | |
| *AaCYP71D10* | AACTTGTGGCGCTGACGA | 60 | Terpenoid |
| | TCTTCGTTCTTGGATTATATTGGC | | |

2. *Amplification of the 10 candidate CYPs through Semi-q-PCR*

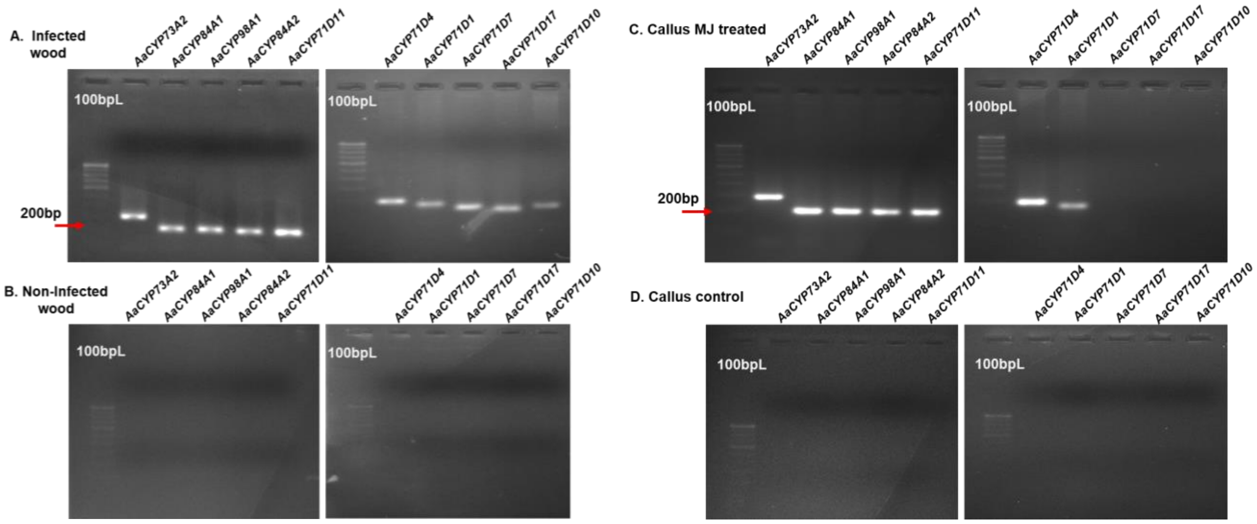


**Fig.** Amplification of 10 CYPs from cDNA converted from A) RNA extracted from infected wood tissue of *A. agallocha*. B) RNA extracted from non-infected wood tissue of *A. agallocha*. C) RNA extracted from callus tissue treated with 0.1mM MeJ for 48hours. D) RNA extracted from non-treated callus tissue.
