## Supplementary figures and images for "Genome-wide investigation of Cytochrome P450 superfamily of *Aquilaria agallocha*: association with terpenoids and phenylpropanoids biosynthesis"

### Additional file 11.pdf

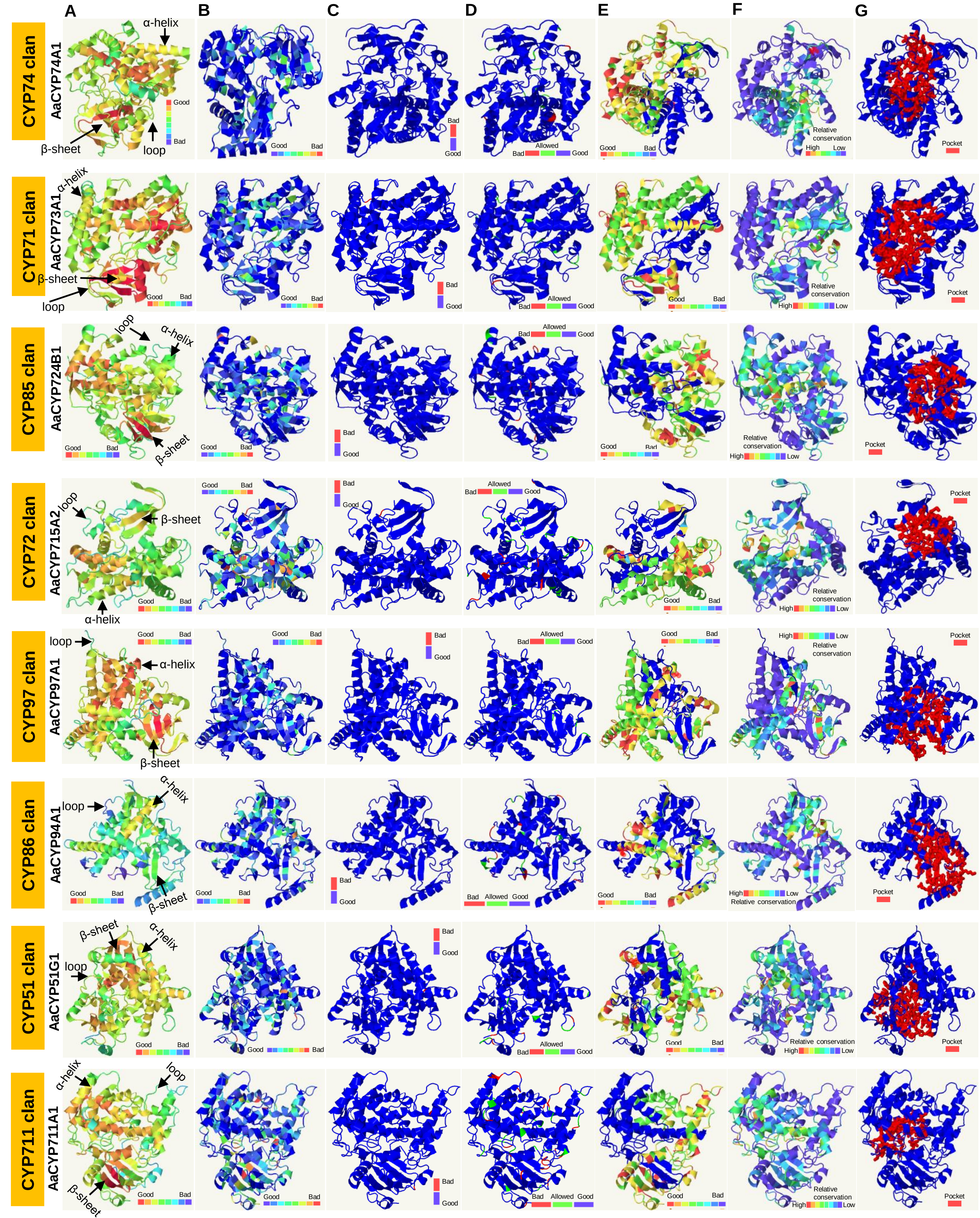

### Additional file 12.pdf

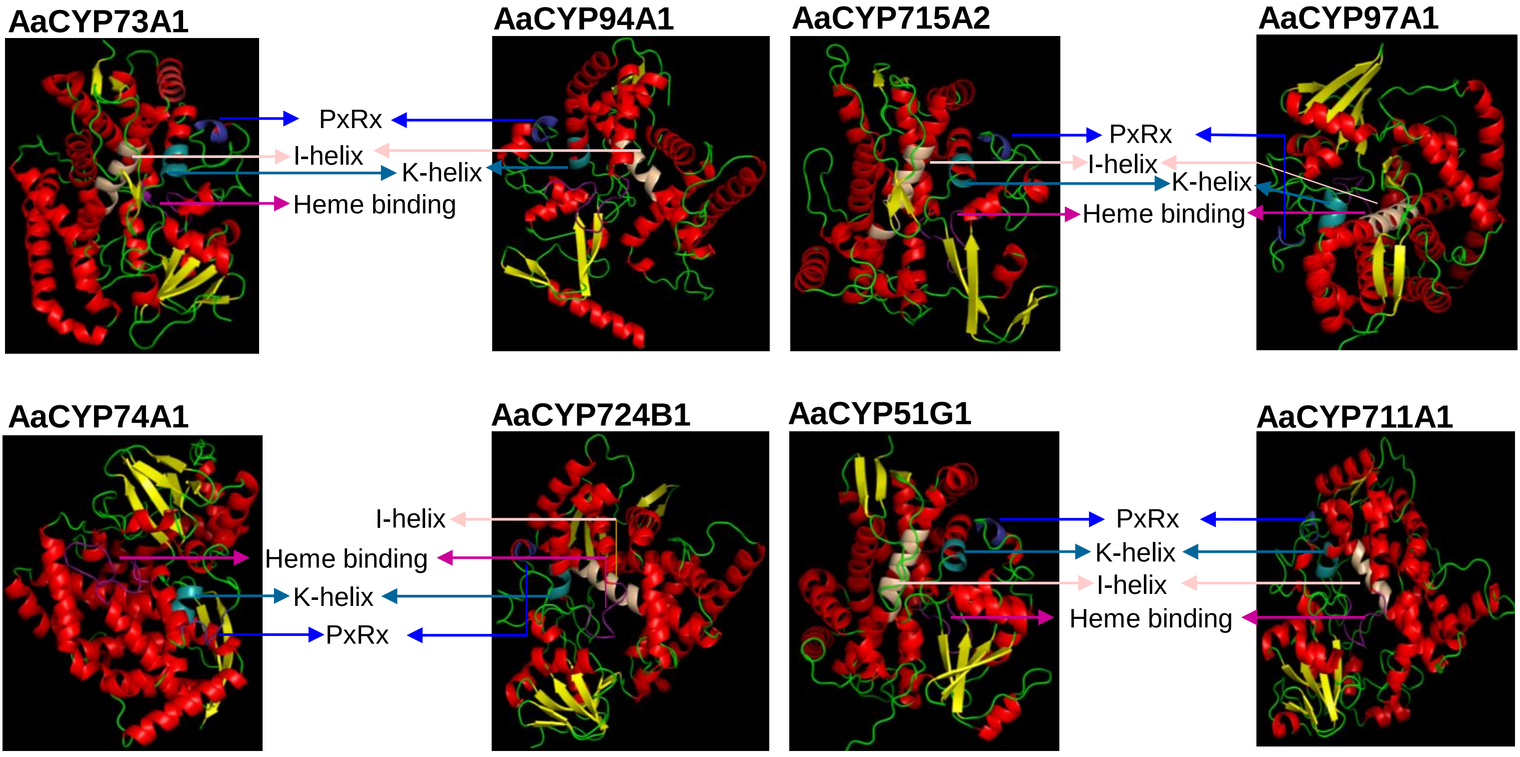
